# Detection of simple and complex *de novo* mutations without, with, or with multiple reference sequences

**DOI:** 10.1101/698910

**Authors:** Kiran V Garimella, Zamin Iqbal, Michael A. Krause, Susana Campino, Mihir Kekre, Eleanor Drury, Dominic Kwiatkowski, Juliana M. Sa, Thomas E. Wellems, Gil McVean

## Abstract

The characterization of *de novo* mutations in regions of high sequence and structural diversity from whole genome sequencing data remains highly challenging. Complex structural variants tend to arise in regions of high repetitiveness and low complexity, challenging both *de novo* assembly, where short-reads do not capture the long-range context required for resolution, and mapping approaches, where improper alignment of reads to a reference genome that is highly diverged from that of the sample can lead to false or partial calls. Long-read technologies can potentially solve such problems but are currently unfeasible to use at scale. Here we present Corticall, a graph-based method that combines the advantages of multiple technologies and prior data sources to detect arbitrary classes of genetic variant. We construct multi-sample, coloured de Bruijn graphs from shortread data for all samples, align long-read-derived haplotypes and multiple reference data sources to restore graph connectivity information, and call variants using graph path-finding algorithms and a model for simultaneous alignment and recombination. We validate and evaluate the approach using extensive simulations and use it to characterize the rate and spectrum of *de novo* mutation events in 119 progeny from four *Plasmodium falciparum* experimental crosses, using long-read data on the parents to inform reconstructions of the progeny and to detect several known and novel non-allelic homologous recombination events.

## 1 Introduction

High genomic diversity within a population can confound variant and particularly *de novo* mutation (DNM) discovery efforts. As a single reference genome cannot capture the range of possible haplotypes, short read aligners assume that new haplotypes are small perturbations to a known canonical reference sequence. Divergent or absent loci violate this assumption, hence reads sampled from them may align incorrectly or not at all(1). This results in many false positives and false negatives in such regions, the combination of which can be sometimes be erroneously interpreted as complex forms of variation. Maps of “genome accessibility” can restrict variant calling to less diverse regions of the genome and reduce such errors(2; 3; 4; 5), but may lead to substantial undiscovered variation. For example, we examined 20 high-coverage *P. falciparum* pedigree samples where one parent is expected to be substantially divergent from the reference. We compared novel sequences present in short-read *de novo* assemblies of progeny versus haplotypes combinatorically produced from multiple reference-based callsets on the same data. Even with strong filtering on the *de novo* assembly data and no filtering on the reference-based callset data, 28% ± 22% (min=0%, max=94%) of novel sequences in the assemblies did not correspond to a reference-based variant call. Most of these unexplained novel “*k*-mers” (length *k* substrings from reads) were found in reads that failed to map or mapped non-uniquely to the reference sequence. This comparison is shown in Figure 1 (see Supplemental Material section S1 for further analysis details).

**Figure 1:**
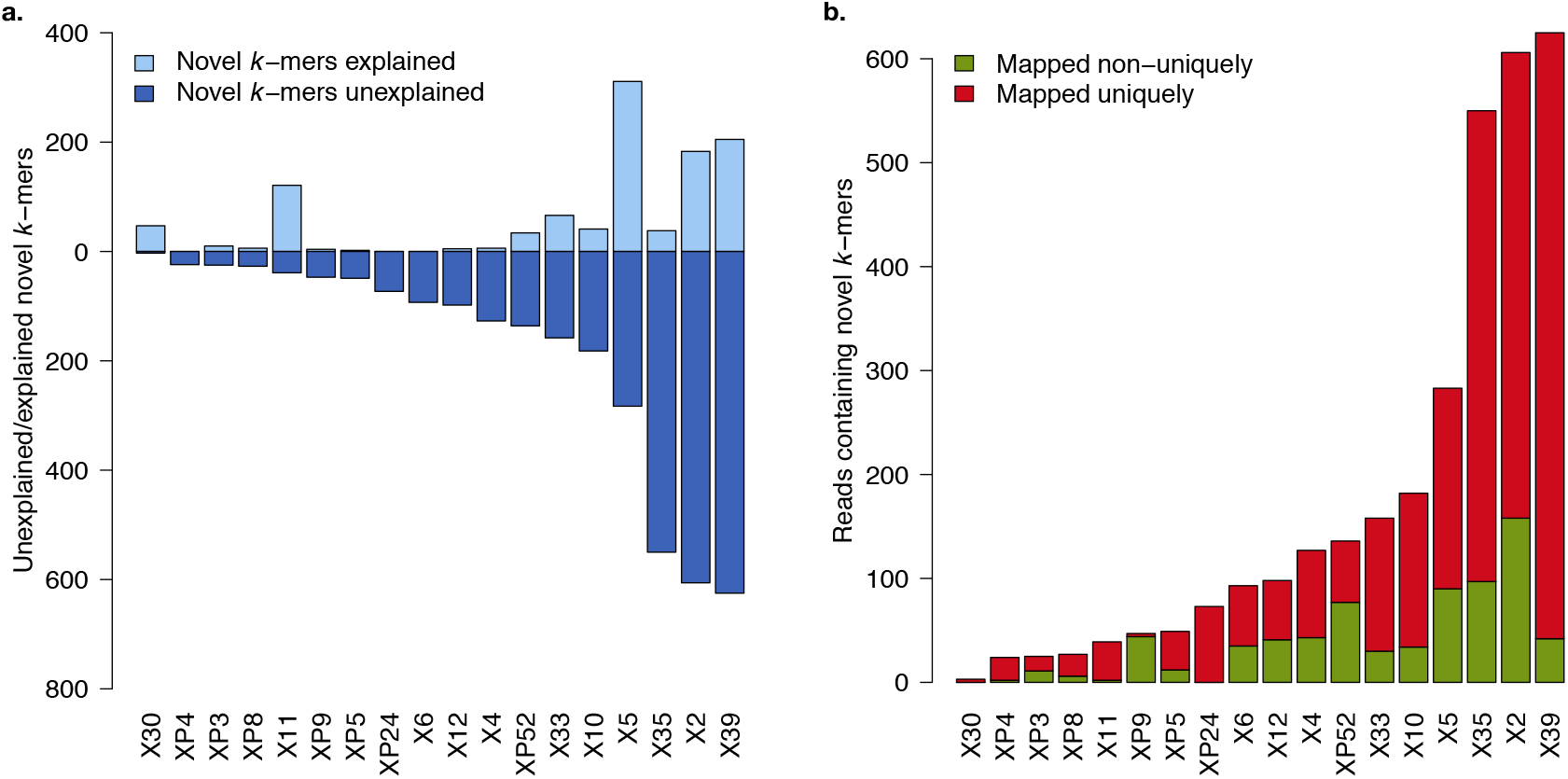
Extent of reference-characterized and uncharacterised novelty among 18 progeny from an experimental cross between 3D7 and HB3 *P. falciparum* isolates (76-bp reads, ~100*x* coverage). (**a**) Novel *k*-mers observed in the reference-based analysis (“explained”, bars above zero-line) versus novel *k*-mers remaining from the reference-free analysis (“unexplained”, bars below zero-line). (**b**) Reads that map uniquely to the reference genome (*MQ >* 0) versus mapping multiple times or not mapping at all (*MQ* = 0), conditioned on the read containing a novel *k*-mer. See Supplemental Material for details.

Of particular concern are *de novo* structural variants (SVs) driven by mutational mechanisms mediated by microhomology and repeat structure(6). Many SVs are predisposed to occur within repetitive loci around the genome. For example, non-allelic homologous recombination (NAHR) can occur between two low copy-number repeats (LCRs), repetitive sequences ranging from several to hundreds of kilobases in length and having > 95% sequence identity between them(7). Non-allelic copies will occasionally be aligned in meiosis and mitosis, with subsequent crossover employing them as the substrate for homologous recombination. Resolution of the misaligned sequences can yield successive insertions, deletions, duplications, inversions, and translocations(8). NAHR in humans has been associated with several genomic disorders (e.g. Charcot-Marie-Tooth 1A, hereditary neuropathy with liability to pressure palsies(9)), and cancer (e.g. hereditary breast/ovarian cancer(10)).

While short read *de novo* assembly may provide a means for overcoming reference bias, the repetitive nature of the SVs precludes the straightforward application of existing tools(11; 12). A typical assembly graph stores genomic subsequences *k*-mers as vertices and sequence overlaps (read-to-read alignments or *k* − 1 substring matches) as edges(13). Repeats longer than the vertex length collapse into a single copy. Differing sequence contexts manifest as multiple edges, which is problematic for assembly as extracting unambiguous contiguous sequence from a graph requires runs of vertices with in-degree and out-degree of 1 (“unitigs”).

For small sample sizes, *de novo* assembly using long read data from third generation sequencing is a viable strategy for overcoming reference bias and assembling through highly repetitive loci(14; 15). However, the high molecular weight gDNA input requirement relative to second-generation sequencing (~10, 000 ng versus ~1 ng) is difficult to satisfy with some samples. Many pathogens grow slowly in culture, requiring several months or even years to expand to sufficient amounts for long-read sequencing. Stromal contamination and high heterogeneity in cancer samples compromises the ability to acquire pure samples of such high mass, and amplification risks PCR replication artifacts masquerading as true *de novo* mutations.

Instead, it may be possible to sequence a small number of samples with long reads to augment a larger, short read data set. For a typical assembly from short read data (e.g. 76 bp reads, > 20*x* coverage), sequencing is expected to recover nearly^*^ every *k*-mer in the genome(16), even if the reads do not provide sufficient genomic context to navigate through repetitive regions. That context can be provided by aligning long haplotypes to the short read graph, annotating edge choices, and following these choices when traversing the graph(17). Importantly, these long haplotypes need not be from the sample itself; recent common ancestry among samples leads to extensive sharing of variation that can be used to guide assembly in related samples. By demanding that the short read genome graph is immutable (after initial construction and removal or correction of likely sequencing errors), the process of long haplotype alignment cannot add any new vertices, only provide connectivity information through existing vertices. This naturally constrains the alignments to informing connectivity in regions of high (but not necessarily perfect) homology between the long read and short read samples. Finally, by aligning multiple data sets to the graph (many long-read datasets, paired-end reads from the sample itself, etc.), we can assemble through recombination breakpoints by transitioning between annotation sets. In essence, rather than using existing tools to improve accuracy of long-read assemblies with short reads(18; 19; 20; 21), we improve the connectivity of short-read assemblies with long reads.

Here, we demonstrate that a small panel of long-read samples could be used to significantly improve *de novo* mutation calling efforts in short read samples. We show that by threading longread-derived haplotypes through short-read graphs, contig lengths spanning putative DNM sites increases dramatically — often by an order of magnitude. We further elucidate a model of simultaneous alignment and “contig painting” wherein progeny contigs are probabilistically aligned to multiple candidate parental haplotypes and recombination between any haplotype and any site is permitted. This model provides a uniform approach to typing simple (e.g. single nucleotide variants — “SNVs”, small insertions/deletions — “indels”, multi-nucleotide variants — “MNVs”) and complex (e.g. large indels or rearrangements manifesting as multiple successive breakends — “BND”) variants.

After testing in simulated datasets, we apply our software to progeny from four experimental crosses of the etiological agent of malaria, *Plasmodium falciparum*. We specifically chose to examine the *P. falciparum* data as the parasite’s genome is of intermediate size (~23Mb, small enough that PacBio sequencing of all parental genomes was feasible, and large enough to force several software design choices that would enable scaling to larger genomes and datasets), notoriously repetitive (*AT* content ~ 80%), haploid (allowing phase between proximate structural variant breakends to be assumed and avoiding the complexity of diploid assembly), and known to possess pathogenically relevant NAHR activity with validated examples. We discover DNMs of all mutational classes in core genomic regions at rates consistent with previous literature, and observe statistically significant enrichment of variants in accessory regions of the genome. Furthermore, we rediscover a known NAHR event and redefine the boundaries beyond those previously established. We also discover new NAHR events, nearly all of which occur within or proximate to antigenic genes. Finally, we examine the transformation of our model from a *de novo* mutation calling framework to a general variant calling framework, showing that the iterative addition of draft reference sequence data yields stepwise improvements in variant discovery. These tools are implemented in a novel graph-based, connectivity-aware variant caller, Corticall.

## 2 RESULTS

Our DNM discovery approach consists of three steps. First, *de novo* assembly, based on multicolor linked de Bruijn graphs, or LdBGs, is used to store and link adjacent *k*-mers for each sample. These assemblies are error-cleaned-that is, low frequency *k*-mers likely to be the result of sequencing errors are removed from the graphs. Unlike error-correction, error-cleaning does not add new (and potentially unobserved) sequence to the graph. Second, trusted “novel” *k*mers are identified; sequences unique to the individual progeny, indicative of DNMs, that are unlikely to arise from error or contamination. Finally, novel *k*-mer-spanning contigs are aligned to reconstructed sequences in the parents, identifying the nature of the event that generated the DNM. Figure 2 depicts these steps, detailed in the Supplemental Material and summarized below.

**Figure 2:**
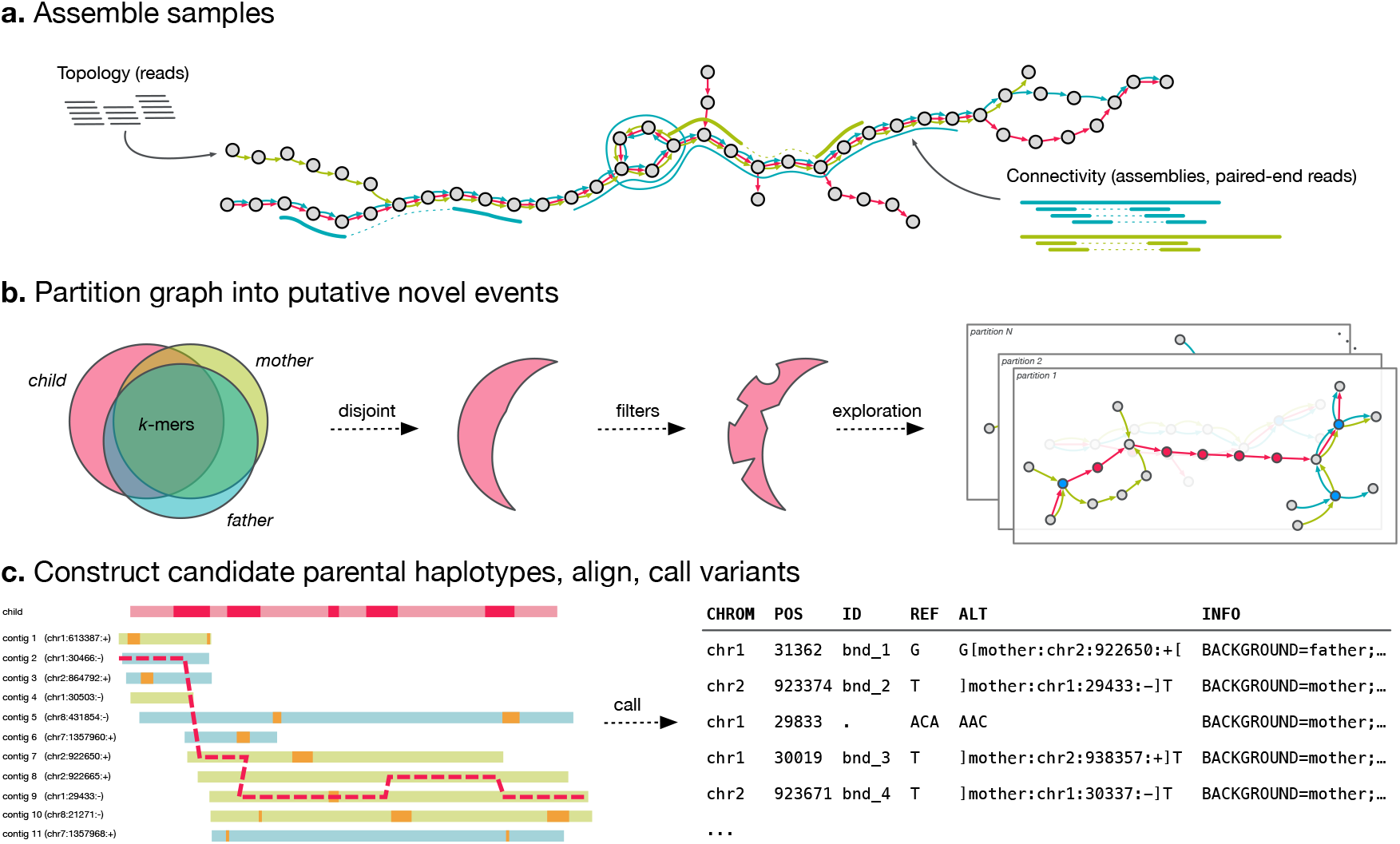
Overview of the Corticall algorithm. (**a**) Samples are assembled into a multi-color linked de Bruijn graph (LdBG). Short, accurate reads are used to determine graph topology. Longer sequences derived from paired-end reads or from draft/finished assemblies are thread through the graph, providing information on connectivity to overcome repeats, but not adding novel *k*-mers. (**b**) Novel *k*-mers-sequences present in the progeny and absent in the parents-are filtered and then used to signal the presence of putative *de novo* mutations. Subgraphs around such events are extracted, forming a set of variant candidates. (**c**) Regions flanking novel *k*-mers are assembled to reveal candidate parental haplotypes. The progeny’s contig is probabilistically aligned to the set of candidate parental contigs, allowing for mismatches, indels, and (potentially non-allelic) recombination. The resulting alignment thus specifies parental background and (if reference sequences are available) coordinate information. Variants (SNVs/MNVs/indels/translocation breakends/etc.) within the novel *k*-mer regions are returned as likely *de novo* mutations.

### 2.1 Connectivity preserved in multi-color linked de Bruijn graphs (LdBGs)

We have previously reported on multi-sample and multi-color de Bruijn graphs (dBG) for straightforward reference-free genome comparison between multiple samples(22) and linked de Bruijn graphs (LdBG) for improved assembly via read-to-graph and reference-to-graph alignment annotations(17). Briefly, an LdBG is a multigraph(23) representation of multiple genomes that preserves “stackability” (easy comparison of multiple samples via inner joins by *k*-mer of per-sample coverage and edge information) and connectivity information inherent in reads and/or long input haplotypes. As illustrated in Figure 2a, input reads are decomposed into *k*-mers and stored as graph vertices. Each sample is assigned a unique identifier (or “color”). Colored edges are placed between vertices representing *k* − 1 overlaps with another *k*-mer in the same sample. Reads and/or haplotype data (e.g. alternate reference assemblies) are then aligned to the graph (once per color) by trivial lookups of shared *k*-mers. Discrepancies between the sequence and the graph manifest as missing *k*-mers, correctable by traversing the graph between the gap boundaries, or truncating the alignment if the correction attempt fails. At junctions (vertices with in-degree or out-degree greater than 1), the edge consistent with the aligned sequence is recorded in an auxiliary file. All junctions spanned by an alignment are annotated with relevant link information, ensuring traversal can begin anywhere in the graph and still have access to complete navigation data. During traversal, we collect links in the order they are encountered, assigning each link an “age” reflecting the number of vertices traversed since collected, and using the oldest link to specify junction choices. If a conflict arises between multiple oldest links, we halt traversal.

### 2.2 Novel *k*-mers are signposts for DNMs

We build upon this genome comparison framework by first identifying regions of the joint pedigree graph (an LdBG containing sequence data for parents, progeny, and optional reference sequences) to explore for potential DNMs. As such mutations are by definition present in the progeny and absent in the parents, *k*-mers spanning these events would also be expected to be exclusive to the progeny.

An accurate list of novel *k*-mers serves both as an indicator of DNM presence around the graph and a measure of how many mutational events are available for discovery. However, iteration over the graph and selection of putative novel *k*-mers (those with 0 coverage in the parents and > 0 coverage in the progeny) will yield a set enriched for sequencing errors and other artifacts that obscure the small fraction of *k*-mers arising from genuine DNMs. We apply multiple filters to remove such artifacts (specifically, contamination; graph tips; low-complexity sequence; “orphans“, sequence found in the progeny but with no edges to parental sequences; low-coverage *k*-mers; “unanchored” *k*-mers, *k*-mers in branches that have no unique alignment in any provided genome; and *k*-mers shared by other progeny, see Supplemental Material section S2.4 for details). We verified these filters by examining novelty in simulated *P. falciparum* crosses and a real trio for which we obtained PacBio sequencing on both parents and progeny.

### 2.3 Contigs spanning novel *k*-mers contain putative *de novo* events

Next, we “partition” the graph into subgraphs, grouping novel *k*-mers into separate bins based on their proximity to one another within the graph. This is illustrated in Figure 2b. Each partition may harbor one or more DNMs, but DNMs are not split across multiple partitions. At each novel *k*-mer, we walk along the progeny’s color in the pedigree graph, exploring outwards and constructing the longest possible contigs. To maximize contig length (and thus increase our sensitivity to complex variation), we employ two strategies. First, links derived from haplotype alignments (e.g. draft references, paired-end reads, etc.) are used to disambiguate junction choices. Second, as DNMs will typically yield a succession of novel *k*-mers in a graph, and as the previous filtering step will have removed most artifacts, we walk past junctions when one (and only one) of the outgoing edges at a novel *k*-mer connects to another novel *k*-mer. This procedure, which we have termed “novel *k*-mer aggregation”, ensures that proximate novel *k*-mers are considered together, useful for large structural variants that may manifest as a series of nearby, but non-adjacent, runs of novel *k*-mers.

### 2.4 Assembling adjacent parental contigs for event decoding

We then construct parental sequences that constitute the candidate haplotypic background(s) for a DNM. At each parentally-shared *k*-mer in a partition, we initiate a contig assembly in the parents. The presence of novel *k*-mers in the partition may lead to gaps in the parental contigs not automatically filled by this assembly step. We close these gaps via depth-first searches (DFS) between bordering *k*-mers. To prevent a combinatoric explosion of considered paths, we limit our explorations to depths of 1, 000 bp by default. For gaps we fail to close in this manner, we assemble flanking boundaries up to a maximum of 500 bp.

Each contig is given a label specifying the parental background from which it was reconstructed and a unique index. If draft/finished reference sequence data is available, we additionally attach coordinate information by aligning each parental contig to the associated draft reference sequence via a built-in version of bwa mem(24)^†^.

### 2.5 “Mosaic” alignment reveals simple and complex mutations

To identify mutations, determine parental background and assign genomic coordinates, we apply a pair-HMM to simultaneously align and phase progeny contigs over candidate parental haplotypes. This model, originally used to study evolutionary relationships in a set of highly diverse antigenic genes from the *P. falciparum var* gene family(25), combines the probabilistic models for sequence alignment(26) and the detection of recombination events(27). Recast in a structural variant framework, it enables simultaneous discovery of both simple/complex mutations in a panel of sequences that are not pre-aligned to one another. As our model permits recombination between any site and any candidate parental haplotype, it also enables the detection of non-allelic events, such as NAHR.

Briefly, the method is as follows. Consider a query sequence (the contig in the progeny) and a set of *N* source sequences (contigs in both parents, partially or completely spanning the target sequence). Our goal is to describe the target sequence as a set of match/mismatch, insertion, deletion, and recombination operations on the source sequences. We choose the starting point in the source sequence uniformly across all sites in the source sequences, beginning in the match or insert states with some probability. At each position, there exists the probability of jumping to any target sequence and any position via recombination. The maximum likelihood alignment (and trajectory through the target panel sequence space) is obtained using the Viterbi algorithm. Variant calls are obtained by examining the traceback path and identifying differences with respect to the query sequence. This process is depicted in Figure 2c. Further mathematical specification is available in the Supplemental Material section S2.7.

A simple set of post-processing filters are applied to keep false discovery rates low. For all mutational types, we reject events containing less than 5 novel *k*-mers. We additionally require NAHR events to satisfy one of two conditions: (1) multiple breakends are detected within a single contig; (2) single breakends are detected within 2, 000 bp of breakends satisfying (1).

### 2.6 Simulation: novel k-mer detection and increased contig lengths

To evaluate our ability to correctly detect DNMs in assembly data, we generated an *in silico* pedigree, simulating full-length (23 Mb) genomes for 1, 000 *P. falciparum* progeny and incorporating a wide range of *de novo* events for later evaluation. Annotated draft reference sequences constructed for two *P. falciparum* isolates (HB3 and DD2, see Supplemental Material section S3) were used as parental genomes. We computed *k*-mer-based homology maps per sister chromatid and modeled crossovers per chromosome based on empirical rates(28), keeping track of the relocated members of each parent’s *var* gene repertoire. We then added simple and complex DNMs, simulating small (1 − 100 bp), intermediate (101 − 500), and large (501 − 1, 000 bp) events, and placing them randomly throughout the genome. In addition to simulating SNVs, MNVs/indels with random sequence, and inversions, special care was taken to simulate variants arising from repeat expansion and contraction by searching for existing repetitive regions in the genome and adding or subtracting repeat units. NAHR events were simulated by recombining members of the progeny’s *var* gene repertoire after meiotic recombination. Assuming a low *de novo* mutation rate, 3 random events were simulated per progeny.

For each progeny’s genome, we simulated 76 bp paired-end reads with an insert size distribution of 250 ± 50 bp, stochastic coverage of 100*X*, and a sequencing error rate of 0.5% (*∼ Q*23).These values were comparable to existing data on the HB3xDD2 cross(29; 30; 3). We constructed joint pedigree graphs using existing Illumina data for the HB3xDD2 parents along with our simulated reads for the progeny, applying the assembly procedure detailed in S4.5 (initial assembly at *k* = 47, error cleaning, and paired-end read and draft reference threading), and extracted novel *k*-mers according to the procedure in S2.4.

We first evaluated our novel *k*-mer detection procedure on these simulated datasets. Figure 3a summarizes our detection of true and false novel *k*-mers. We were able to detect 90.0% ± 22.7% of expected novel *k*-mers per sample. Novel *k*-mers that we failed to detect were typically low-complexity or repetitive sequences (generated *de novo* by the mutation process but also occurring elsewhere in the genome). For these events, a *k*-mer size of 47 bp was insufficient to resolve the sequences as novel.

**Figure 3:**
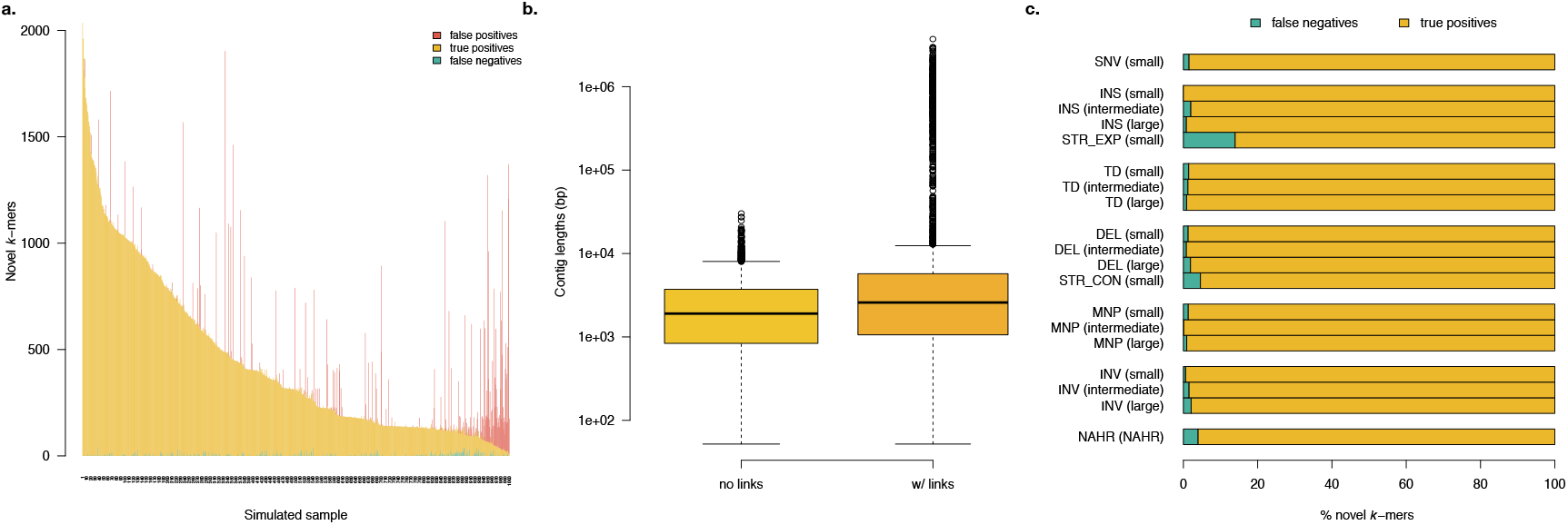
Simulation-based evaluation of novel *k*-mer detection and subsequent reassembly quality for contigs spanning novel *k*-mers in error-containing short read data. (**a**) Number of *k*-mers in the progeny correctly identified as novel (true positives), undetected (false negatives), and misidentified as novel (false positives). (**b**) Contig lengths resulting from reconstruction strategies without and with link information. (**c**) For all simulated alleles, the fraction assembled completely (i.e. wholly contained within a single contig) and incompletely (i.e. only partially reconstructed).

Next, we studied our novel *k*-mer-spanning contig reconstructions. Contigs spanning novel *k*-mers were assembled with three different procedures; resulting lengths were calculated and aggregated across all 1, 000 simulated samples and are presented in Figure 3b. Without link information, median contig N50 lengths (minimum contig length required to cover 50% of the genome) were 4, 186 bp. This increased by ~ 71% to a median of 7, 161 bp with the addition of links from paired-end reads and parental assembly links (Figure 3b).

Finally, we sought to understand the relationship between missed novel *k*-mers and the type of variant event from which they arose, summarized in Figure 3c. Across all variant types, 97.8% ± 3.1% of novel *k*-mers generated by mutational events are detected. The bottom three performers are short tandem repeat (STR) contractions, STR expansions, and NAHR events, where the percentage of novel *k*-mers detected are 86.1%, 95.4%, and 96.0% respectively. This is to be expected; all three mutational classes are manipulations of repetitive sequence, the expansion/contraction/recombination of which would be plausibly expected to generate *k*-mers already present in other repeats in the genome.

### 2.7 Simulation: mutation detection and evaluation

We applied our software to the simulated set of 1, 000 HB3xDD2 progeny, measuring performance using *F*_1_ scores for variants. The aggregated results for all simulated samples are shown in Table 1.

**Table 1:** Variant classification and *F*_1_ score in 1, 000 simulated *P. falciparum* genomes.

| Type | Length (bp) | N | Without link information |  | With link information |  |
| --- | --- | --- | --- | --- | --- | --- |
| | | | $F_1$ | $F_1$ (w/ phase) | $F_1$ | $F_1$ (w/ phase) |
| SNVs | 0 | 627 | 0.85 | 0.85 | 0.86 | 0.86 |
| Insertions |  |  |  |  |  |  |
| random sequence | 1-100 | 29 | 0.81 | 0.74 | 0.83 | 0.76 |
|  | 101-500 | 111 | 0.99 | 0.91 | 0.99 | 0.91 |
|  | 501-1000 | 145 | 0.98 | 0.90 | 0.99 | 0.91 |
| STR expansions | (variable) | 323 | 0.91 | 0.83 | 0.92 | 0.85 |
| tandem duplications | (variable) | 287 | 0.69 | 0.64 | 0.70 | 0.65 |
| Deletions |  |  |  |  |  |  |
| random sequence | 1-100 | 29 | 0.75 | 0.70 | 0.81 | 0.73 |
|  | 101-500 | 125 | 0.84 | 0.80 | 0.84 | 0.81 |
|  | 501-1000 | 444 | 0.53 | 0.52 | 0.54 | 0.52 |
| STR contractions | (variable) | 301 | 0.91 | 0.87 | 0.92 | 0.90 |
| MNVs |  |  |  |  |  |  |
| random sequence | 1-100 | 32 | 0.85 | 0.79 | 0.93 | 0.82 |
|  | 101-500 | 106 | 0.73 | 0.72 | 0.76 | 0.75 |
|  | 501-1000 | 135 | 0.77 | 0.76 | 0.79 | 0.78 |
| inversions | 1-100 | 29 | 0.86 | 0.84 | 0.91 | 0.82 |
|  | 101-500 | 116 | 0.88 | 0.88 | 0.89 | 0.88 |
|  | 501-1000 | 137 | 0.88 | 0.88 | 0.89 | 0.88 |
| NAHRs |  |  |  |  |  |  |
| single breakpoints | - | 1142 | 0.63 | 0.63 | 0.70 | 0.70 |
| all breakpoints | - | 270 | 0.60 | 0.60 | 0.66 | 0.66 |

<sup>a</sup> For NAHR events, partial (complete) reconstruction indicates recovery of any (all) of the simulated breakends.

Overall, we found that greater than 90% of detected novel *k*-mers are assignable to variant events, and more than 97% of simulated variants are identified (either partially or completely reconstructed). This changes very little with assembly mode as, aside from some light filtering, the absence or presence of link information does not alter the detection of novel *k*-mers. Instead, it simply alters the number of contigs into which a variant assembles. For complete reconstruction of each variant event, *F*_1_ uniformly increases between the link-uninformed and link-informed reconstruction as link information provides a means to overcome repetitive regions of the assembly. This is particularly valuable for DNMs in repetitive elements: STR expansions, STR contractions, and tandem duplications. For these variant classes, link information conferred a ~ 10%, ~ 10%, and ~ 14% increase in *F*_1_ respectively.

We measured calling performance on breakends, and further our ability to group proximate breakends into single NAHR events. While both reconstructions are generally able to detect the presence of a breakend, the LdBG reconstructions show marked improvement in event characterization. This permit multiple breakends to be observed on a single contig, enabling detection and assignment of all breakends within a simulated NAHR event to a single call.

### 2.8 Core, accessory, and repetitive region DNM detection in *P. falciparum*

To characterize the number and type of DNMs occurring in the genome of the malaria parasite *P. falciparum*, we applied our software to data from four *P. falciparum* experimental crosses: 3D7xHB3(31), HB3xDD2(29), 7G8xGB4(32), and 803xGB4(33). In addition to the 3D7 canonical reference genome(34), we generated high-quality draft reference assemblies for the remaining five parental genomes (HB3, DD2, 7G8, GB4, and 803) using PacBio RSII sequencing (Supplemental Table 3). We obtained Illumina data for all samples in the experimental crosses (Supplemental Table 4) and generated McCortex assemblies at *k* = 47. After contaminant and outlier removal, we called DNMs in 119 progeny. These calls are summarized in Figure 4.

**Figure 4:**
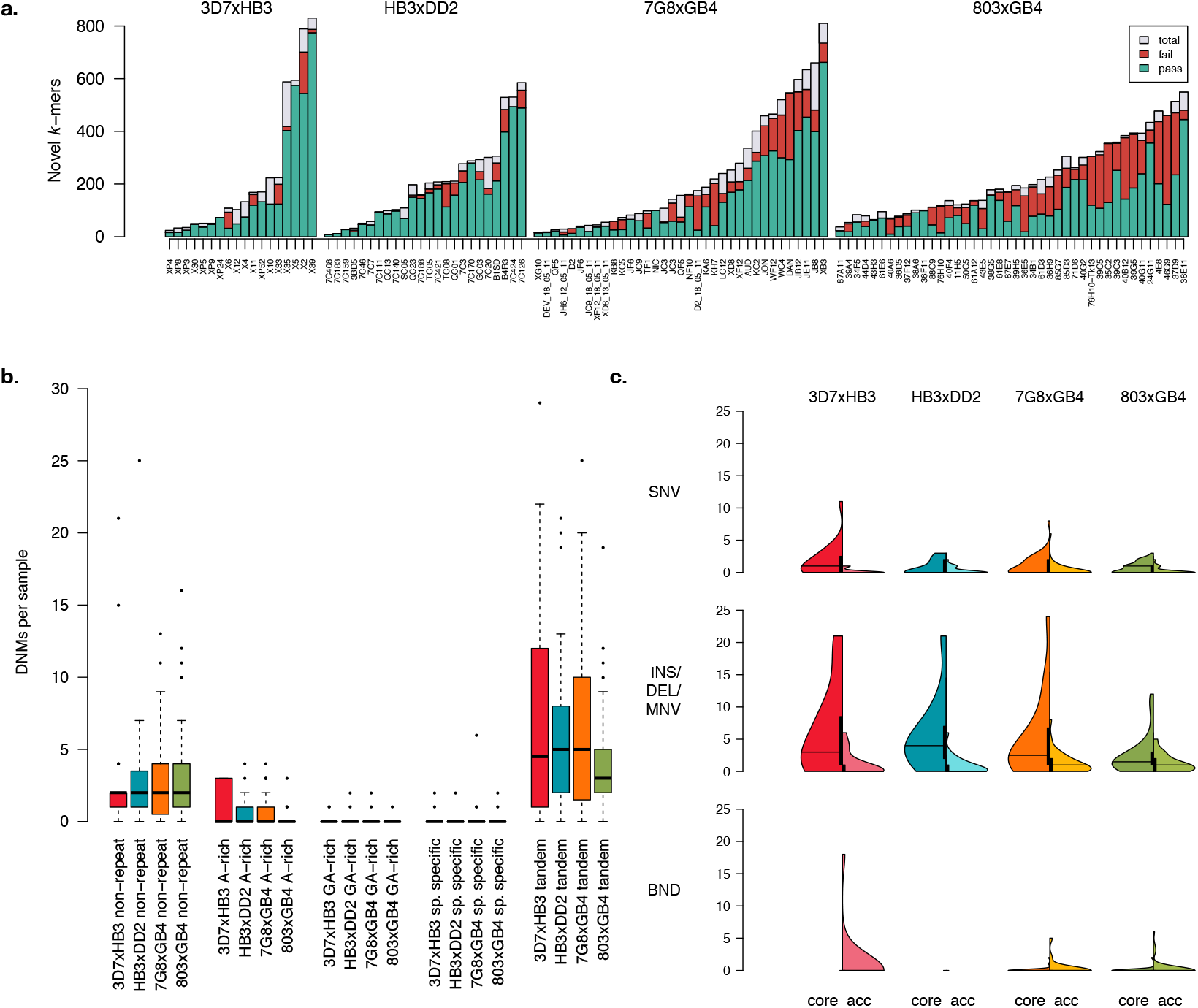
Per-sample DNM discovery metrics in 119 *P. falciparum* progeny. (**a**) Novel *k*-mers per cross and sample (grey bars). For those contained within successfully assembled variants, *k*mers in variant passing filters are shown in green; the rest are shown in red. (**b**) Per cross DNM sample distributions for mutations appearing in repetitive regions of the respective parental genomes. (**c**) Violin plots showing DNM sample distributions per cross, split by those in core genomic regions (left) and accessory regions (right).

Across samples, we assigned putative variant calls to (89 ± 11)% of novel *k*-mers (Figure 4a). Their impact was greatest in the 803xGB4 cross, where the 803 and GB4 draft reference sequences have comparatively poorer assembly qualities of *Q*28 and *Q*23, respectively (compared to *Q*28 and *Q*29 for 3D7 and HB3). After filtering, we detected a total of 972 *de novo* mutations (163 SNVs, 348 insertions, 322 deletions, 19 MNVs, 7 NAHR events, and 113 incompletely assembled events).

The average per sample DNM count is low, with short indels (~ 5.47 per sample) outnumbering SNVs (~ 1.29 per sample). We applied the RepeatMasker(35) software to annotate repetitive genomic sequences and Spine(36) to annotate accessory genomic regions (sequences private to each parasite isolate, typically encompassing subtelomeric/hypervariable regions) across all parental genomes. We then inspected variant locations with respect to these annotations (Figure 4b-c). Aggregated across all samples and crosses, we found a 3-fold enrichment of mutations occurring in repetitive genomic regions, ~ 90% of which fell within tandem duplications. Mutations were enriched in the accessory (~ 2 Mb) versus core (~ 21 Mb) genomic compartments (SNVs: *p* = 1.5 *×* 10^−7^; INS/DEL/MNV: *p <* 2.2 *×* 10^−16^; BND: *p <* 2.2 *×* 10^−16^; based on chi-squared tests accounting for indel lengths and number of novel *k*-mers appearing in NAHR events, see Supplemental Material). We observed similar per-sample mutation distributions across samples.

We computed per-sample per-nucleotide mutation rates across all four crosses. Additionally, as DNMs can continue to accumulate in each parasite during the *in vitro* intraerythrocytic lifecycle, we computed mutational rates per nucleotide and generation. However, culture time and lifecycle time for cross progeny was not always known. Assuming a culture time of 52 days between initial cloning and sequencing (the average of the documented culture times for the 3D7xHB3 and HB3xDD2 cross progeny), and a mitotic generation time of 48 hours(37), Per-nucleotide mutational rates are presented in Table 2. These rates are broadly consistent across crosses and compartments, and with previous estimates based on parasite clone trees(38; 39).

**Table 2:** Mutation rates per cross, sample, mutational class, and genome compartment

|  | 3D7xHB3 | HB3xDD2 | 7G8xGB4 | 803xGB4 |
| --- | --- | --- | --- | --- |
| Culture time (days)<sup>a</sup> | 47 | 57 | 52<sup>b</sup> | 52<sup>b</sup> |
| Lifecycle time (hours)<sup>c</sup> | 48 | 48 | 48 | 48 |
| Progeny | 18 | 24 | 35 | 42 |
| Genome length (bp)<sup>d</sup> |  |  |  |  |
| core | 20810915 | 21052828 | 21325706 | 21303692 |
| (accessory) | (1860495) | (1603876) | (2368812) | (2389390) |
| Total variants<sup>e</sup> |  |  |  |  |
| SNVs | 32 (3) | 21 (7) | 43 (14) | 33 (10) |
| Indels | 114 (19) | 141 (21) | 188 (46) | 106 (54) |
| NAHRs | 0 (2) | 0 (0) | 0 (2) | 0 (3) |
| Rate ($sample^{-1} bp^{-1}$)<sup>e</sup> | | | | |
| SNVs | 8.1e-09 (8.5e-08) | 3.9e-08 (1.7e-07) | 5.3e-08 (1.6e-07) | 3.5e-08 (9.5e-08) |
| Indels | 1.7e-06 (4.0e-06) | 1.8e-06 (1.4e-06) | 1.4e-06 (4.6e-06) | 8.1e-07 (3.3e-06) |
| NAHRs | 0 (6.3e-06) | 0 (0) | 0 (7.9e-07) | 0 (9.4e-07) |
| Rate ($sample^{-1} bp^{-1} gen^{-1}$)<sup>e</sup> | | | | |
| SNVs | 3.4e-09 (3.6e-09) | 1.4e-09 (6.1e-09) | 2.0e-09 (5.9e-09) | 1.4e-09 (3.7e-09) |
| Indels | 1.2e-08 (2.3e-08) | 9.4e-09 (1.8e-08) | 8.9e-09 (1.9e-08) | 4.3e-09 (1.9e-08) |
| NAHRs | 0 (2.4e-09) | 0 (0) | 0 (8.5e-10) | 0 (1.1e-09) |

<sup>a</sup> Culture time estimates from Claessens *et al.* 2014.<sup>b</sup> Assumed as mean of 3D7xHB3 and HB3xDD2 culture times.<sup>c</sup> Assumed from Trampuz *et al.* 2003.<sup>d</sup> Averaged lengths of core and accessory regions for each parent.<sup>e</sup> Core (accessory) rates shown outside (inside) parentheses.

**Table 3:** Nearest genes to NAHR breakends

| Gene | 3D7 ortholog | Encodes | Function |
| --- | --- | --- | --- |
| Pf3D7_0100100 | - | PfEMP1 | immune evasion |
| Pf3D7_0223500 | - | PfEMP1 | immune evasion |
| Pf3D7_0700100 | - | PfEMP1 | immune evasion |
| Pf3D7_1100100 | - | PfEMP1 | immune evasion |
| Pf7G8_010005000 | unknown | hypothetical protein, conserved | unknown |
| Pf803_070005000 | Pf3D7_0100100 | PfEMP1 | immune evasion |
| PfGB4_010005200 | unknown | hypothetical protein, conserved | unknown |
| PfGB4_030005000 | unknown | hypothetical protein, conserved | unknown |
| PfGB4_050039900 | Pf3D7_0700200 | RIF | variant surface antigen |
| PfGB4_080041200 | Pf3D7_0100100 | PfEMP1 | immune evasion |
| PfGB4_110005000 | Pf3D7_0223500 | PfEMP1 | immune evasion |
| PfGB4_120005100 | unknown | hypothetical protein, conserved | unknown |

### 2.9 Hypothesis-free discovery of NAHR events to basepair resolution

To detect NAHR events, we grouped proximate breakend calls and applied three filtration criteria: (1) events must contain 20 or more novel *k*-mers, (2) consist of 3 or more breakends, and (3) at least one contig must link distal genomic loci within the same contig. We detected 7 NAHR events in total after applying, depicted in Figure 5. All occurred in subtelomeric accessory regions of the genomes. To determine genes nearest to each NAHR breakend, we transferred existing 3D7 gene models and performed *ab initio* gene prediction on each parental genome via the Companion(41) annotation server. All but four of the genes closest to a breakend were related to antigenic gene families and immune evasion.

**Figure 5:**
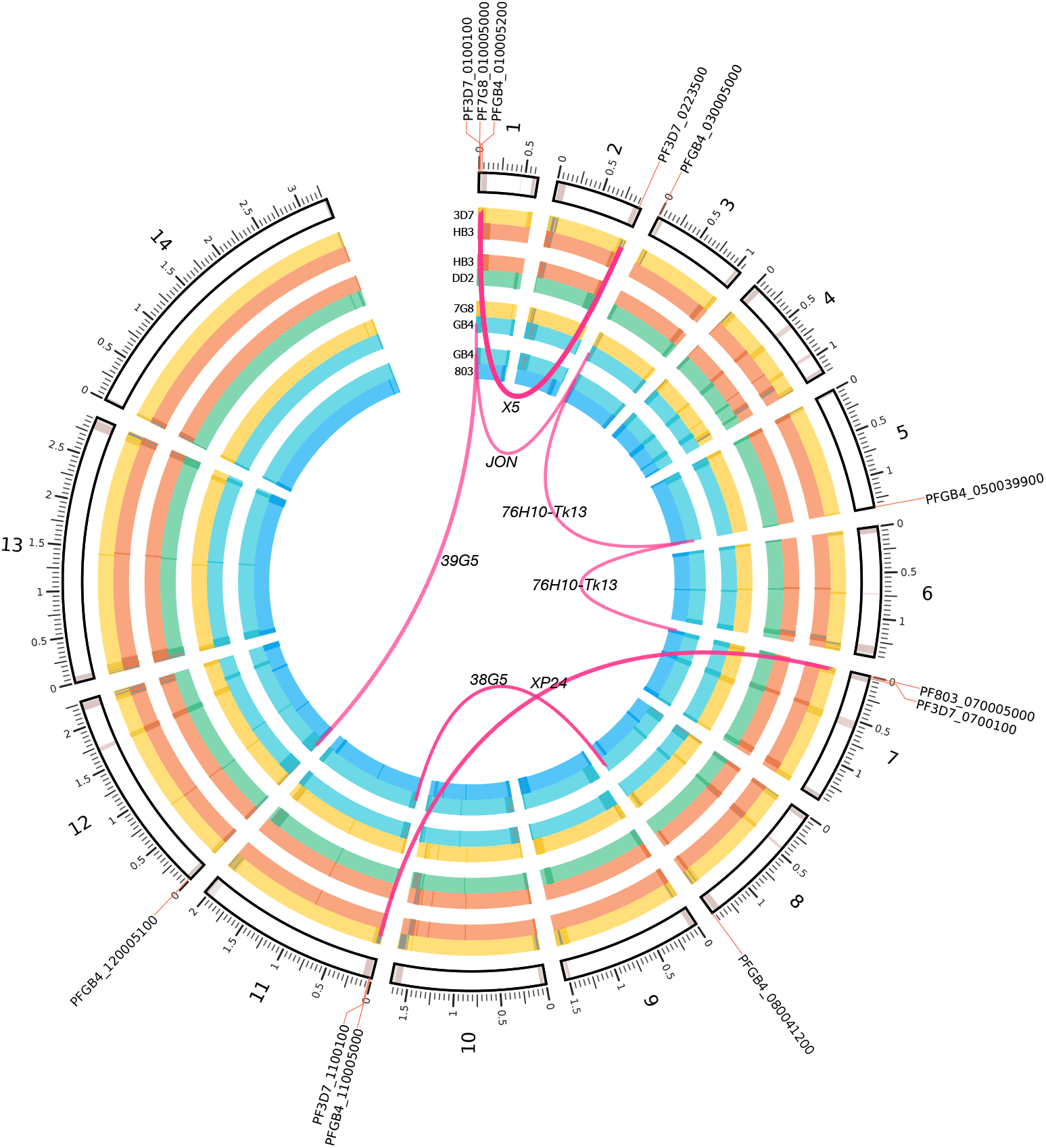
Circos(40) plot of NAHR events detected in all 119 samples across four*P. falciparum* experimental crosses. Parental genomes for each cross are depicted in the inner grouped circular tracks. Beézier curves depict each translocation event, with termini indicating the parent(s) of origin and a label at the apex of the curve identifying the sample in which it was found. Closest gene names annotated on outer circumference. Dark bands indicate accessory regions determined by the Spine(36) software, except in the outer ideogram, which is based on alignability maps for the canonical 3D7 reference genome(3).

Previous work on NAHR events in *P. falciparum* – based on observations of apparent translocations of *var* gene sequences, and limited by inadequate reference sequences for parasites other than 3D7 – have only reported NAHR events within the exon 1 of *var*-gene family members(38; 39; 42; 43; 44; 45; 46). As we enforce no *a priori* hypothesis on which loci are likely to harbor such recombinations, the discovered events in our dataset extend beyond *var* exon 1. While the events still occur in the subtelomeric regions of the genome (within which many other genes related to immune evasion reside), five of the twelve genes proximate to the NAHR breakends were not *var* genes. A single event occurred near a gene from the *rif* gene family.

Beyond identifying new NAHR events outside of the usual *var*-gene repertoire, we were also able to clarify the extent of previously observed events. Figure 6 depicts three of the detected NAHR events. In Figure 6a, our calls recapitulate previously reported rearrangements (breakends 5-9) within the long exons of Pf3D7 0100100 and Pf3D7 0223500^‡^(46). Flanking these known breakends are a number of mutations that have not been previously reported, including an additional series of breakends upstream of each of the *var* genes (1 and 2), two MNVs (3 and 4), and an SNV within the coding region of the antigenic gene on chromosome 2. In panel b, a novel NAHR event is shown with a recombination path that weaves in and out of coding regions, touching upon the previously unexamined exon 2. The recombination path within the novel event in panel c (within a sample in the previously unpublished 803xGB4 cross), remains wholly within the coding sequence.

**Figure 6:**
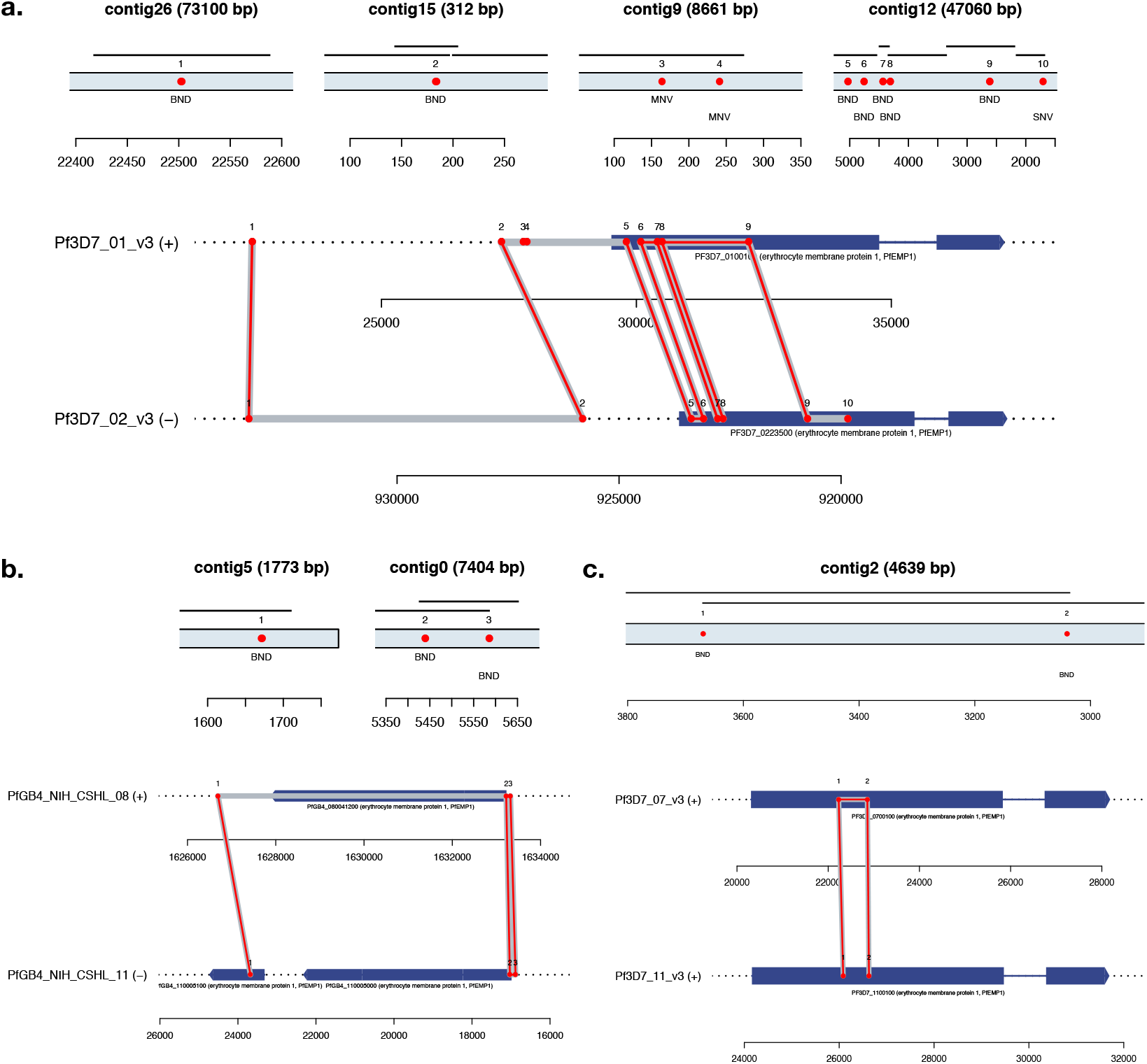
Three of the detected NAHR events in the *P. falciparum* crosses. (**a**) NAHR event involving two *var* genes in 3D7xHB3 progeny X5 (Pf3D7 0100100 on chr1, Pf3D7 0223500 on chr2). Top: LdBG contigs spanning mutation (dBG contig shown as thin black line for comparison). Called mutations shown along contig as red points. Bottom: mutations from LdBG contigs in genomic context shown in red. Gene models shown in dark blue (thick lines: exonic sequence; thin lines: intronic sequence). Inferred recombination path shown in gray. (**b**) NAHR event in 803xGB4 sample 38G5 (PfGB4 080041200 on chr8, PfGB4 11005100 and PfGB4 11005000 on chr11). (**c**) NAHR event in 3D7xHB3 sample XP24 (Pf3D7 0700100 on chr7 and Pf3D7 1100100 on chr11).g

### 2.10 Variant calling with cumulatively expanding reference set

Exploring beyond comparisons of progeny-to-progenitor genomes, we hypothesized that genomic novelty present in a sample but not placeable on the background of an evolutionarily distant reference sequence would be better elucidated through the simultaneous use of multiple reference sequences. We obtained Illumina data and constructed a PacBio draft assembly of an 803xGB4 progeny (36F11). From the 36F11 data, we extracted *k*-mers that were novel with respect to the 3D7 reference genome, further filtering these *k*-mers based on presence in the counterpart 36F11 clone assembly, thus constructing a conservative *k*-mer list that flags true variation in the 36F11 parasite. We used this list to seed variant calls, increasing the number of reference sequences provided with each callset.

Figure 7 depicts the calling results on 36F11 with the cumulative addition of 3D7, HB3, 7G8, DD2, GB4, and 803 reference sequences. As novel *k*-mers are computed with respect to 3D7, calls at these *k*-mers can only be homozygous-variant. As additional reference sequences are added, variants are described against a new background sequence. However, many novel *k*-mers tagging variation against 3D7 are no longer considered novel w.r.t. another reference sequence, and their reconstructed sequence for the progeny is perfectly homologous to the additional reference. Thus, as more reference sequences are added, apparent variation against 3D7 is re-described as homozygous-variant (hashed bars) or homozygous-reference to a sequence other than 3D7 (solid bars). When using all six reference sequences, our ability to characterize apparent novelty to 3D7 grows from 40% to 95%.

**Figure 7:**
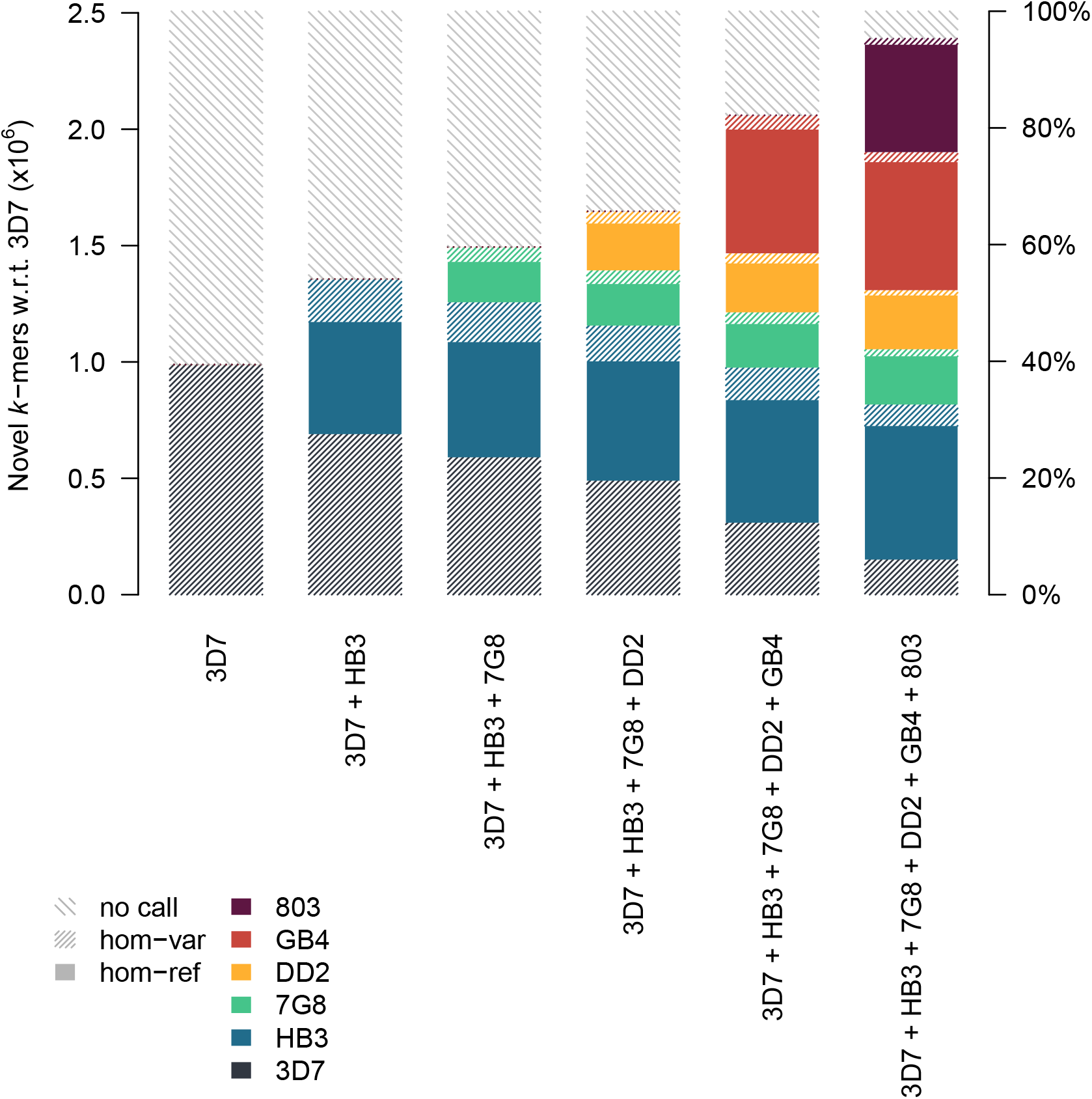
Calls tagged by 36F11 *k*-mers novel with respect to 3D7, redescribed against combinations of other reference sequences. Stacked bars represent fraction of novel *k*-mers linked to homozygous-reference (hom-ref) and homozygous-variant (hom-var) calls, or *k*-mers where no call could be made. Colors represent the specific haplotypic background the call was placed on (if a call can be equally described on multiple backgrounds, one is chosen at random).

## 3 DISCUSSION

We have presented a graph-based *de novo* mutation calling method, available through our software Corticall, that is capable of discovering simple and complex variants in pedigrees and experimental crosses without bias towards a reference sequence. Our approach leverages long-haplotype data derived from any source (existing finished genomes, draft assemblies from thirdgeneration sequencing, targeted sequencing of specific loci, etc.) to improve the assemblies of other short-read datasets. These long-haplotype samples need not be from the same sample. Short-read data is used to establish graph topology, while long-haplotype data is aligned to the graph but constrained to specify connectivity information only. Sequencing errors (and possibly mutations) are always adjudicated in favor of the existing graph, thus no new sequence is added, only navigation information. This approach opens the opportunity for multiple longread datasets to be used to improve the connectivity of many more short-read assemblies.

Corticall can leverage many finished or draft reference-quality datasets, seamlessly transitioning between connectivity information sets during assembly. This affords a powerful approach to the hypothesis-free study of *de novo* mutations. As many of these events occur in repetitive or genetically diverse regions of the genome, the use of multiple reference sequences during assembly helps to provide access to so-called genomic “dark matter” - loci underserved by pure short-read *de novo* assembly or a single canonical reference.

Corticall assembles variants, not genomes, and keeps false discovery rates low by only inspecting regions of the genome harboring novel *k*-mers. By combining local, multi-sample assembly with a simultaneous alignment/recombination model, we are able to detect a wide variety of mutational types with a single, consistent framework. Additionally, tracking the number of novel *k*-mers explained by each variant call provides a useful metric for determining the completeness of the final callset.

In the *P. falciparum* crosses, we detected SNVs at rates broadly consistent with previous work, and indels at more than four times the SNV rate. We detected new NAHR events, all in subtelomeric regions of the genome that are not represented in the canonical reference. For previously discovered NAHR events, we are able to find additional breakends in nearby noncoding regions, establishing a more complete picture of non-allelic recombination behavior in these pathogens. Much of the *de novo* mutational spectrum appears in accessory regions. These compartments are diverse in the population precisely because they typically harbor clinically relevant genes underlying drug resistance or immune escape functionality. The mapping-free, reference-agnostic approach espoused by Corticall thus enables the detection of this clinically-relevant variation, and removes the requirement for determining the appropriate genome reference for mapping and analysis.

The fixed record size structure of Cortex graphs used with Corticall enables storage in an ordered, randomly accessible manner, thus keeping memory requirements low as the entire graph need not be loaded into memory in order to be inspected. Pre-determining the novel *k*mers to inspect, along with intelligent caching to prevent redundant lookups when assembling multiple samples over shared *k*-mers, reduces disk accesses. As a result, Corticall is able to scale to genomes of any size. This may provide a valuable approach to the study of Mendelian disease in large pedigrees or tumor/normal pairs (wherein the normal can be considered as the parent of the tumor samples).

Corticall has several limitations, addressable by future work. While Corticall need not load an entire graph into memory to perform variant calling, the genome assembly software upon which it relies *does* require that the entire graph be stored in RAM as it is being constructed. Thus, even though the variant calling step on human data can be done in as little as 1 Gb of RAM, the initial *de novo* assembly step still requires hundreds of gigabytes of memory to execute. Recent approaches to streaming graph construction(47) and/or succinct de Bruijn graphs(48; 49) may well address this limitation.

Additionally, our use of long haplotype data is restricted to sequences that have been substantially error-corrected. Typically *k*-mer sizes used in de Bruijn graph-based short read assemblies (e.g. *k* = 31 − 96) are still too high for the long, error-prone reads generated by third-generation sequencers. However, lowering the *k*-mer size of the short read assemblies to a length more likely to result in a perfect match on the long read data (e.g. *k* = 11) would result in too many junctions from homologous sequences in the graph. Our current approach to error-correcting long reads against the graph requires that the path through the existing graph contain no junctions, and would thus be impaired by setting the *k*-mer size to low. A more computationally expensive read-to-graph alignment procedure could remedy this limitation.

As third-generation sequencing continues to mature, the construction of additional draft reference genomes will become more accessible. The utility of this data extends beyond pure *de novo* assembly for constructing new reference sequences, or for elucidating structural variation in single samples. Strategic choices as to which samples to sequence with long reads can enable simple and complex variant discovery in a much larger cohort while simultaneously keeping costs low, provided that variant calling methods are capable of leveraging such information. Corticall is a step forward in this direction, presenting a uniform approach to variant discovery and typing that combines assembly, alignment, recombination models, and third-generation reference sequence panels. Such approaches will assist in overcoming bias to a single canonical reference sequence and enable a more complete description of variation in diverse populations.

**URLs.** Corticall, part of the CortexJDK package: https://github.com/mcveanlab/CortexJDK.

## 4 METHODS

Methods and associated references are available in the Supplemental Material for this manuscript.

Accession codes. PacBio and Illumina sequence data have been deposited at the European Nucleotide Archive (ENA) (see Supplemental Material for accession numbers).

## ACKNOWLEDGEMENTS

The authors thank Eric Antoniou, Sara Goodwin, Michael Schatz, and the CSHL PacBio sequencing service; Winni Kretzschmar and Karl Johan Westrin for helpful code improvements reducing the memory usage of the Tesserae model; and Isaac Turner, Patrick Albers, Jerome Kelleher, and Marcus Tutert for helpful technical discussions and manuscript review. This work was supported by grants from the Wellcome Trust (numbers 090532/Z/09/Z and 100956/Z/13/Z) and the Li Ka Shing Foundation (to G.M.). K.V.G. was supported by Wellcome Trust Research Studentship award (097310/Z/11/Z). Z.I. was funded by a Wellcome Trust/Royal Society Sir Henry Dale Fellowship (102541/Z/13/Z).

## AUTHOR CONTRIBUTIONS

K.V.G. developed algorithms and pipelines for identifying mutations in sequence assembly graphs. M.A.K., S.C., M.K., E.D., and J.S. oversaw the culturing of malaria parasites and subsequent DNA extraction for PacBio sequencing. D.K. and T.W. provided lab support for the parasite culturing and access to Illumina data on all four *P. falciparum* crosses. Z.I. and G.M. provided access to critical resources. K.V.G., Z.I., and G.M. wrote the manuscript.

## Footnotes

* barring systematic sequencing errors and ultra low-complexity sequences that fail to amplify.

† bwa mem Java bindings developed by Pierre Lindenbaum, https://github.com/lindenb/jbwa

‡ PFA0005w and PFB1055c in older nomenclature

