## Supplemental Material for "Detection of simple and complex *de novo* mutations without, with, or with multiple reference sequences"

*

#### Introduction

Here, we provide further detail on the motivation, methods, data acquisition, processing, and analysis steps underlying the results presented in this work. Software, manuscript source, and additional resources (e.g. genome assemblies, machine-readable text files with paths to data downloads) are available at the following locations:

| Purpose | URL | License |
|---|---|---|
| Genome assembly | `https://github.com/mcveanlab/mccortex` | MIT |
| Variant assembly | `https://github.com/mcveanlab/CortexJDK` | Apache 2.0 |
| Additional resources | `https://github.com/mcveanlab/CortexJDK/tree/master/manuscript` | - |

#### S1 Missing novelty in reference-based scans for *de novo* mutations

Our main manuscript presents an analysis wherein genomic novelty in several *P. falciparum* isolates is quantified using two separate methods and compared. This analysis is detailed below.

Consider a scenario in which genomic sequence sampled from an individual is absent or sufficiently disparate from a canonical reference genome. Many sequenced reads may fail to align to the reference correctly. Variants spanned by those reads may thus be rendered undiscoverable by reference-based approaches to variant detection. However, quantifying this missingness — the number of *de novo* mutations (DNMs) in a sample that remain to been discovered — is not straightforward.

We hypothesized that this missingness could be quantified by comparing a reference-*free* DNM novel sequence set (i.e. one constructed by performing *de novo* assembly on all members of the pedigree and emitting haplotype sequences unique to the children) to a reference-*based* novel sequence set (i.e. one constructed by aligning pedigree reads to a reference sequence, identifying mutations, and emitting haplotype sequences implied by those mutations). In an ideal scenario where divergence between samples and the reference is very low, the reference-free and reference-based analyses should be equivalent. With real data, the lack of equivalence can provide insight into how much novelty within a sample remains to be identified.

We performed such an analysis on *P. falciparum* progeny from the 3D7xHB3 experimental cross (20 progeny, 76-bp reads, $\sim$100x coverage, see Table S5 for further detail). The analysis workflow follows and is additionally diagrammed for clarity in Figure S1.

##### S1.1 Reference-free novel sequences

DNMs can be considered generators of novel genomic sequence — sequence that is present in an individual's genome but absent from their parents. Rather than counting mutational events explicitly, we instead count sequences unique to the genome of the child. We used the McCortex software(1) to split sequenced reads into fixed-length subsequences of length $k = 47$ ("$k$-mers"), aggressively removed sequencing errors and other artifacts using McCortex's $k$-mer-frequency-based cleaning and additional filters described in section S2.4, and compute the disjoint between child's $k$-mers and parental $k$-mers (Figure S1a-c). This provides a conservative reference- and alignment-free method of computing the *total* amount of novelty in a child's genome.

##### S1.2 Reference-based novel sequences

We aligned parental and progeny reads to the canonical *P. falciparum* reference of the 3D7 isolate using `bwa mem`(2), called SNVs and small indels using HaplotypeCaller(3) (GATK v4.0.2.1 with the `-ploidy 1` setting) and larger indels and structural variants using Delly(4) (v0.7.8). We did not apply any filtration steps in an effort to generate a callset with maximum sensitivity.

To generate $k$-mers that could be compared with the reference-free analysis, we sought to permute the reference sequence with mutations from the callset and emit $k$-mers spanning the mutations. Given the high false discovery rate of the unfiltered callset, we could not rely on a single haplotype with all variants incorporated. Instead, we generated nearly all possible variant haplotypes within small genomic windows. At each putative mutation, we collected the previous 50-bp of reference sequence and a maximum of 10 variants within the immediately following 50-bp window. We combinatorically constructed all haplotypes from every possible subset of

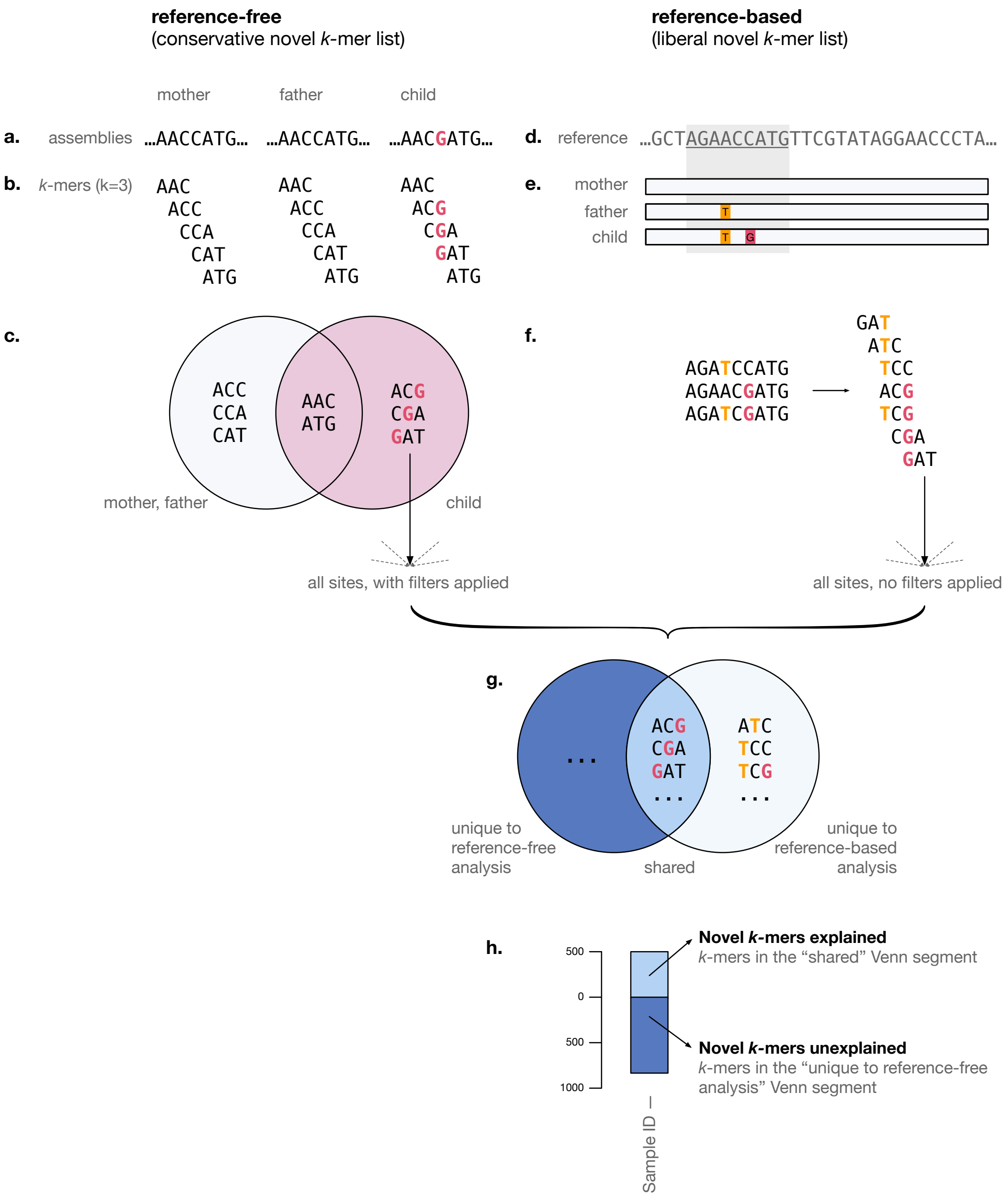

**Figure S1. Procedure for comparing reference-free and reference-based scans for genomic novelty.** *Reference-free pipeline:* (**a**) Reads are assembled *de novo* and error-cleaned into a multi-color de Bruijn graphs using the McCortex software. (**b**) Error-cleaned assemblies are converted to lists of parental and child $k$-mers ($k = 3$ depicted). (**c**) Lists of $k$-mers are compared, and novel $k$-mers (the child's disjoint) are extracted and subjected to additional filtering. *Reference-based pipeline:* (**d**) Reads are aligned to a reference sequence and variants are called using maximum sensitivity settings and no application of filters. (**e**) Windows around putative variants are chosen. (**f**) Variants within window are combinatorically used generate all possible haplotypes, then distilled to $k$-mers of the same size used in the reference-free pipeline. No further filtering is applied. *Comparison:* (**g**) All $k$-mers from other sites in the genome are aggregated. The Venn diagram of $k$-mers from the reference-free and reference-based analyses is computed. (**h**) Shared $k$-mers reflect those that are present in the reference-free analysis and recapitulated by the reference-based analysis. Those unique to the reference-free analysis reflect those that could not be recapitulated by the reference-based analysis. The $k$-mers unique to the reference-based analysis are discarded.

variants collected. Assuming all sites are biallelic, this effectively imposed a theoretical limit of $2^{10} = 1024$ haplotypes from any given site. Additionally, as Delly emits a consensus haplotype sequence for structural variants precisely identified, we included this sequence in the list of haplotypes to be processed. We then emitted $k$-mers from these haplotypes, and advanced to the next unprocessed variant. These steps are depicted in Figure S1d-f.

In our *P. falciparum* data, the restriction of processing the first 10 variants seen within each 50-bp window affected $0.26\% \pm 0.06\%$ of windows processed per sample ($23277 \pm 4325$ windows per sample, $62 \pm 19$ windows reduced per sample).

##### S1.3 Comparison

Finally, we examined the Venn diagram of reference-free and reference-based novel $k$-mer lists, identifying shared $k$-mers (present in both analyses) and reference-free-unique $k$-mers (novel $k$-mers not recapitulated by the reference-based analysis). Novel $k$-mers unique to the reference-based analysis were discarded as these largely originate from the combinatoric haplotype construction procedure and were not true genomic sequences. Figure S1g-h show these final comparison steps.

#### S2 Methods for DNM discovery based on linked genome assembly graphs

##### S2.1 Overview

Our DNM calling strategy is based on identifying mutational motifs in a “multi-color linked de Bruijn graph”, or LdBG(5; 6). This can be decomposed into three steps. First, we construct LdBGs from short-read and long-haplotype datasets. Second, for each so-called “novel” $k$-mer (those unique to a child and absent from its parents), we assemble a child contig and one or more parental contigs containing $k$-mers shared with the child contig. Finally, we perform probabilistic all-to-all alignment allowing for recombination, attempting to describe the child’s sequence as a series of match, insertion, deletion, and recombination operations on a panel of candidate parental sequences. Decoding the traceback of the probabilistic alignment yields variant calls. Further detail on each step is provided below.

##### S2.2 Construction of the linked de Bruijn graph

Briefly, a de Bruijn graph for sample $c$ is formulated as a set of vertices and edges, $\mathcal{G}_c = \{\mathcal{V}_c, \mathcal{E}_c\}$. Vertices $\mathcal{V}_c$ are input sequences are broken into fixed length substrings of length $k$ (“$k$-mers”) with unit stride, and edges $\mathcal{E}_c$ encode $k-1$ overlaps of adjacent vertices. Each record is recorded as three columns: a $k$-mer sequence, its coverage, and its incoming/outgoing edges. $N$ sample graphs constructed at identical $k$ can be “stacked” by performing a full (outer) join on $k$-mer sequences, each sample $c$’s coverage and edge information simply being recorded as two additional columns in each $k$-mer record. Stacking facilitates easy comparison of the graphs of $N$ samples and formally yields a union graph $\mathcal{G} = \bigcup_{c=1}^{N} \mathcal{G}_c$. This formulation encodes relationships between two adjacent $k$-mers (the $i$-th and $(i+1)$-th $k$-mers in a sequence, as well as the $(i-1)$-th and $i$-th), but relationships between non-adjacent $k$-mers are lost. Thus, even if an input sequence spans a repeat when a single $k$-mer does not, the connectivity information inherent in the sequence is not retained. We restore this connectivity by trivially aligning input sequences $\mathcal{R}_{c,d}$ from dataset $d$ to graph $\mathcal{G}_c$. The addition of new vertices to the graph during alignment is disallowed; the process merely amounts to lookups of shared $k$-mers between the input sequence and the graph, and bridging gaps over sequence differences with simple walks on $\mathcal{G}_c$. For all junctions (vertices with in-degree or out-degree $> 1$) spanned by an input sequence, we record the series of disambiguating edge choices (referred to as “links”), exhaustively annotating all participating junctions with relevant navigation information. We refer to this composite data structure (graph and links) as a linked de Bruijn graph, $\mathcal{G}_c = \{\mathcal{V}_c, \mathcal{E}_c, \bigcup_{d=1}^{D} \mathcal{L}_{c,d}\}$, where $\mathcal{L}_{c,d}$ is a sparse set of links on graph color $c$ derived from sequence dataset $d$.

##### S2.3 Using links during LdBG navigation

By exhaustively annotating all spanned junctions with links, we ensure that traversal initiated anywhere in the graph has access to complete link information. Upon initiating a walk at vertex $v_c$, we collect each link we encounter. At a junction, we consult our list and extract the oldest link (i.e. the link that was obtained earliest in the traversal), as this link establishes the greatest context as to location in the genome. If there are multiple links with the same age that disagree as to the next junction choice, we halt traversal.

##### S2.4 Identification and filtration of novel *k*-mers

In a multi-color de Bruijn graph representing parents and children from a pedigree or an experimental cross, the locations of most DNMs will be signalled by the presence of novel *k*-mers: sequences unique to a child's genome and absent from both parental genomes. The set of novel *k*-mers in a child should also provide an indication as to how much novelty in a genome remains to be explained by some mutational process. As sequencing errors and sample contamination will also contribute to the set of novel *k*-mers, we sought to identify all novel *k*-mers in a child's graph and remove potential errors and contaminants. We identified and developed filters for five common graph or sequence motifs indicative of error:

1. **Contamination**: Contamination presents as a subset of novel *k*-mers that are unique to the sequencing data for a child but are irrelevant to the study at hand. To remove these sequences, each entry in the initial set of putative novel *k*-mers was screened for contamination via BLAST(7). We rejected any *k*-mer with a match of any quality to an organism other than the species under study. To account for mutations present in our contaminants but absent in the BLAST database, we used the contaminating *k*-mers as starting points for depth-first searches (DFS) in our graphs, exploring the child's graph until it rejoins a parent's graph, and rejecting all *k*-mers along the way.
2. **Graph tips**: Graph tips present as a series of novel *k*-mers that bifurcate from a parental graph but never rejoin. They are typically the result of sequencing errors at the ends of reads, but could also reflect true variation and subsequent coverage drop-out during sequencing. However, in the latter case, such variation tagged by novel *k*-mers would still not be recoverable without further sequencing data to fill in the missing coverage. To remove graph tips, we perform DFS from a putative novel *k*-mer, expecting to rejoin a parental graph on both ends. If exploration on one end connects to a parent and fails on the other end, we reject all child *k*-mers contained in the traversal.
3. **Promiscuously connected sequences**: Low-complexity sequence (or "dust") may manifest as *k*-mers promiscuously connected to many other low-complexity *k*-mers, presenting as a unnavigable graphical tangle. We defined such dust *k*-mers as those having a sum of in-degrees and out-degrees greater than 4. We initiated DFS at such *k*-mers, exploring until we either run out of edges to navigate or rejoin a parental graph, and keeping track of the number of *k*-mers traversed since the last time we observed one of low-complexity. If we reach one of the aforementioned stopping conditions and the distance traversed since the last low-complexity *k*-mer is less than the graph's *k*-mer size, we consider the traversed vertices to be dust and reject all elements.
4. **Highly-compressible sequence**: Additional low-complexity sequences are detected by computing the compression ratio ("CR") of the *k*-mer (gzip-compressed length vs uncompressed length) and removing any putative novel *k*-mer with a CR less than a predefined threshold (by default 0.703 for 47-bp *k*-mers).
5. **Orphans**: Graphical orphans are a series of novel *k*-mers that fail to ever connect to a parental graph. They may include contaminants absent from the BLAST database or reads with unusually high sequencing error. We performed DFS at putative novel *k*-mers, rejecting *k*-mers from traversals that joined one of the parental colors at any time.

We also removed putative novel *k*-mers from consideration based on two additional criteria:

1. **Shared *k*-mers**: Putative novel *k*-mers, while absent from parents, may be shared amongst children. Some of these may reflect recurrent *de novo* mutations, but the overwhelming majority stem from recurrent sequencing errors. We remove *k*-mers shared with other children (omitting clones of a child from consideration).
2. **Low coverage**: A number of putative novel *k*-mers substantially less than the mean coverage of the sample. Such *k*-mers may still permit navigation to flanking regions with coordinates in a parental genome, despite arising from sequencing error. We remove *k*-mers with coverage less than a specified value (by default, 6*x*).

The bulk of sequences captured by these final two filters are likely to be recurrent sequencing error. However, we note that they could also remove a small number of DNMs from our consideration.

##### S2.5 Query sequence assembly

To construct sequences spanning putative variants, we perform contig assembly at each novel *k*-mer on the query sample (e.g. the child). Unless otherwise specified, these assemblies are conducted using McCortex links generated by threading the sample's paired-end read data and the parental assembly data through the query sample's graph(1). Optionally during graph traversal, if we encounter a junction vertex that (1) is itself a novel *k*-mer, (2) cannot be traversed with links, (3) one (and only one) of the outgoing vertices is also a novel *k*-mer, then we assume both novel *k*-mers are part of the same mutational event and extend contig construction through these vertices. As assemblies seeded by proximate novel *k*-mers may result in redundant contigs, we post-process the contig set to remove redundant sequences and those fully contained by other contigs. Finally, if multiple contigs share a novel *k*-mer, we remove all but the contig that contains the largest number of novel *k*-mers. This effectively "partitions" the contig set into those representing distinct mutational events.

##### S2.6 Source sequence assembly

For each query sequence, we build a panel of source sequences to which the query is aligned. At each non-novel *k*-mer in the query sequence, we perform contig assembly on the source samples (e.g. the parents). Unless otherwise specified, these assemblies are conducted using McCortex links generated by threading the sample's paired-end read data and the parental assembly data through the child's graph. During assembly, gaps at the boundaries of mutational events in the query sample may be incompletely assembled due to sequencing error or graph homology. We close these gaps via DFS between gap boundaries. If still not closed, we assemble gap flanks by a maximum of 500 bp. Flanking sequence irrelevant to the query is trimmed by subsetting the source within the boundaries of the earliest and latest *k*-mers shared with the query sequence.

Each source sequence is given a unique label, simply incrementing from first to last. If a reference sequence is specified for the relevant sample in the LdBG, the source sequence is aligned to that reference using `bwa mem` and relabelled with the resulting genomic coordinates. Note that the relabelling step does not alter the source sequence in any way.

##### S2.7 Variant typing by simultaneous alignment to reference genome panels

Two general classes of graphical variant motifs concern us: "bubbles" (SNVs, short indels and inversions, multi-nucleotide polymorphisms), and "breakends" (large indels and inversions, nonallelic homologous recombinations, gene conversions, and allelic recombinations). We address both classes of variants in a single probabilistic framework wherein a novel $k$-mer-spanning contig ("query" sequence) is simultaneously aligned to a panel of candidate haplotypes ("source" sequences). We achieve this by repurposing the Tesserae model (8, supplementary methods), a pair-HMM combining models for global alignment with affine gap penalty (described in 9) and haplotype diversity estimation via recombination(10), to the task of bubble and breakpoint variant typing.

The model (including formal descriptions of the Viterbi, Forward, and Backward algorithms) is fully specified in Zilversmit *et al.* 2013. Briefly, we assume a query sequence arises as an imperfect mosaic of source sequences. For each query and its candidate source sequences (collectively referred to as the "sequence set", $h$), we apply the Viterbi algorithm to find the maximum likelihood path through our pair-HMM. The pair-HMM is specified by a transition matrix and emission matrix, provided below. Terms are defined as follows:

**Table S1.** Definitions and defaults in the Tesserae pair-HMM.

| Term | Definition | Default |
|---|---|---|
| $\delta$ | probability of indel initiation | 0.025 |
| $\epsilon$ | probability of indel extension | 0.75 |
| $\rho$ | probability of recombination | 0.0001 |
| $\pi_M$ | probability of starting in match (M) state | 0.75 |
| $\pi_I$ | probability of starting in insert (I) state | $1-\pi_M$ |
| $\tau$ | probability of termination | 0.001 |
| $\lvert Y\rvert$ | $\Sigma_{k=1}^{n} l_k$ where $l_k$ is the length of sequence $k$ | - |

The transition matrix is given by:

| | $B$ | $M_x$ | $I_x$ | $D_x$ | $M_k$ | $I_k$ | $D_k$ | $T$ |
|---|---|---|---|---|---|---|---|---|
| $B$ | 0 | $\frac{\pi_M}{\lvert Y\rvert}$ | $\frac{\pi_I}{\lvert Y\rvert}$ | 0 | $\frac{\pi_M}{\lvert Y\rvert}$ | $\frac{\pi_I}{\lvert Y\rvert}$ | 0 | 0 |
| $M_x$ | 0 | $1-2\delta-\rho-\tau$ | $\delta$ | $\delta$ | $\frac{\rho\pi_M}{\lvert Y\rvert}$ | $\frac{\rho\pi_I}{\lvert Y\rvert}$ | 0 | $\tau$ |
| $I_x$ | 0 | $1-\epsilon-\rho-\tau$ | $\epsilon$ | 0 | $\frac{\rho\pi_M}{\lvert Y\rvert}$ | $\frac{\rho\pi_I}{\lvert Y\rvert}$ | 0 | $\tau$ |
| $D_x$ | 0 | $1-\epsilon$ | 0 | $\epsilon$ | 0 | 0 | 0 | 0 |
| $M_k$ | 0 | $\frac{\rho\pi_M}{\lvert Y\rvert}$ | $\frac{\rho\pi_I}{\lvert Y\rvert}$ | 0 | $1-2\delta-\rho-\tau$ | $\delta$ | $\delta$ | $\tau$ |
| $I_k$ | 0 | $\frac{\rho\pi_M}{\lvert Y\rvert}$ | $\frac{\rho\pi_I}{\lvert Y\rvert}$ | 0 | $1-\epsilon-\rho-\tau$ | $\epsilon$ | 0 | $\tau$ |
| $D_k$ | 0 | 0 | 0 | 0 | $1-\epsilon$ | 0 | $\epsilon$ | 0 |
| $T$ | 0 | 0 | 0 | 0 | 0 | 0 | 0 | 1 |

The emission probability of a nucleotide emitted from an insertion state is $e(x_i) = 0.2$, while the nucleotide emission matrix $e(x_i, y_j)$ (where $x_i$ is the nucleotide at site $i$ in sequence $x$ and $y_j$ is the nucleotide at site $j$ in sequence $y$) is given by:

$$e(x_i, y_j) = \begin{cases} 0.9 & x_i = y_j \\ 0.05 & \text{if nucleotide transition} \\ 0.025 & \text{if nucleotide transversion} \end{cases}.$$

The trellis diagram for the model is provided in Figure S2.

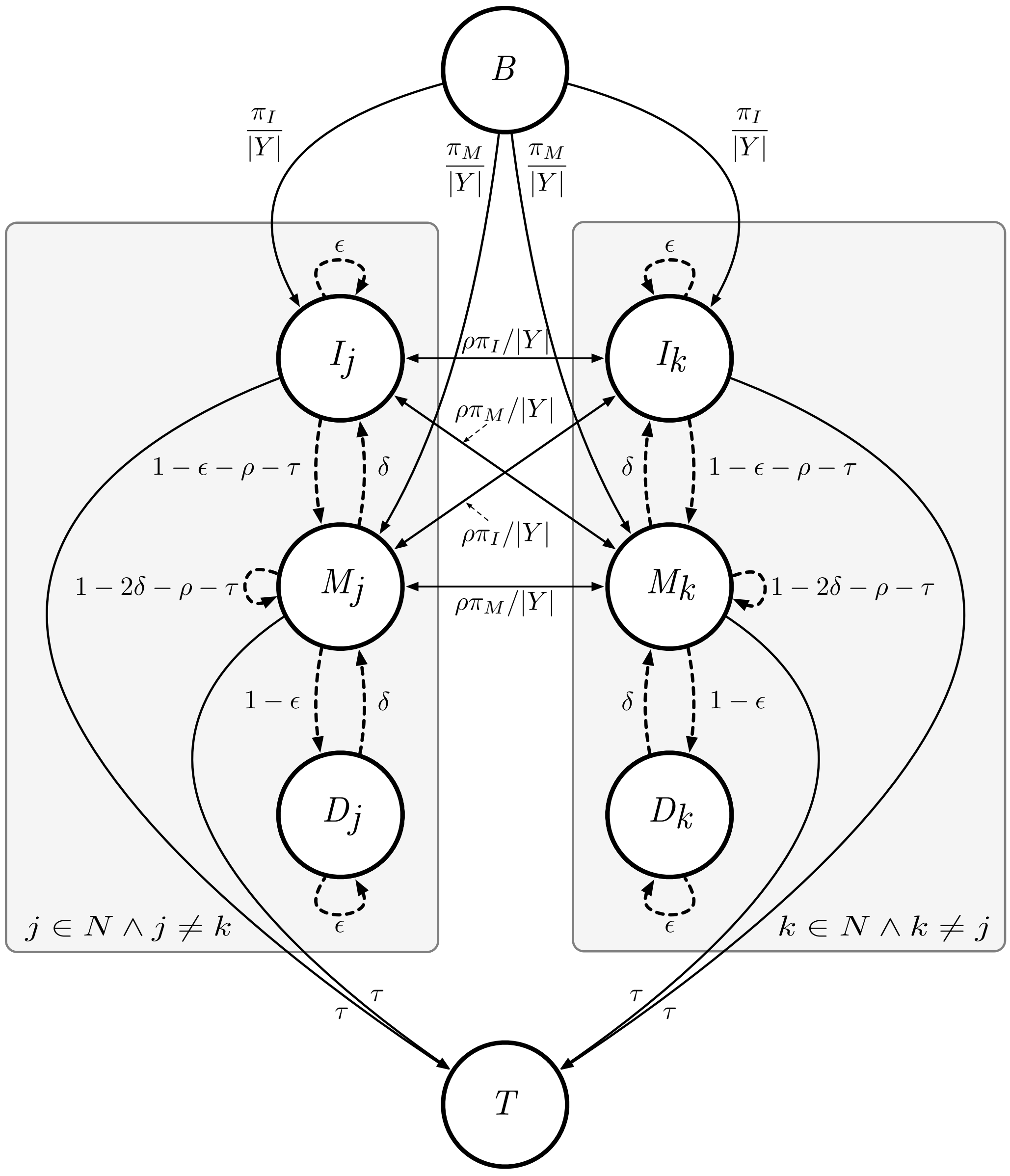

**Figure S2. The Tesserae model trellis diagram for simultaneous alignment with recombination.** Each plate denotes a global alignment model for input query and source sequences. Match, insert, and delete states are connected within single sequences. To permit recombination, all sequences are exhaustively connected through match and insert (but not delete) states. Dashed lines indicate internal (intra-plate) transitions. Solid lines indicate external (inter-plate) transitions.

By keeping pointers through the recursion process of the Viterbi algorithm, we trace back the most probable path through $h$. This path through the resulting traceback matrix can be interpreted as specifying the background haplotype for subsequences and variants in the query sequence against this background. Figure S3 depicts the overall process on a toy example consisting of a short query sequence and two candidate source sequences. The query sequence is constructed by initiating assembly on the LdBG in the query color at one or more novel $k$-mers. The two candidate source haplotypes are constructed by initiating assembly at non-novel $k$-mers found in the query sequence. Redundant contigs are removed (this step is not shown). The sequence set is processed with the Tesserae model, and decoding of the resulting traceback matrix reveals two recombination breakends and an insertion.

```
a.        query   CGAACAGGATGTAGGCGAGATGACGCCATTTATTCTTTTCGTGCATAACAAAACGATAGTAG
        source0   CGAACAGGATCAGGGATAAAACAAATTGATTATTCTTTTCGTGCATAACACGATAGTAG
        source1  GTCATACGACCGTAGGCGAGATGACGCCATTTATTACGGATATTATATTTATATA

b.    query   (0-61) CGAACAGGATGTAGGCGAGATGACGCCATTTATTCTTTTCGTGCATAACAAAACGATAGTAG
                     ||||||||||||||||||||||||||||||||||||||||||||||||^^^||||||||||
   source0    (0-9)  CGAACAGGAT
   source1 (11-34)             GTAGGCGAGATGACGCCATTTATT
   source0 (34-58)                                     CTTTTCGTGCATAAC---ACGATAGTAG

c.                             BND                     BND              INS
```

**Figure S3. Assembly, alignment, and variant decoding process for a toy example.** (**a**) A 61-bp query sequence with three regions of sequence novelty (nucleotides with a red background). To assemble candidate source sequences, *k*-mers in the non-novel regions are used to seed contig traversals on the LdBG in the source colors (in this example, $k = 7$). Sequences with yellow backgrounds indicate the seed *k*-mers for these traversals. For visual clarity, sequences shown here are positioned manually, and source subsequences that are not present in the query sequence are shown in grey. (**b**) All sequences in the sequence set aligned with the Tesserae model, with breakends depicted as linebreaks and variants shown on their haplotypic background. (**c**) Putative mutations spanned by novel *k*-mers are emitted as breakend ("BND") variant calls.

### S3 Data processing: long-read sequencing, assembly, and alignment

#### S3.1 Data availability

PacBio RSII raw read ERA accession numbers, processed assembly download paths, and annotation file download paths are provided below:

**Table S2.** Pacbio read and assembly metadata and download paths.

| source code | secondary sample accession | origin | culturing | sequencing | chemistry | fasta |
|---|---|---|---|---|---|---|
| HB3 | ERS712858 | Honduras | SI | SI | P6C4 | FASTA |
| DD2 | ERS639545 | Vietnam | SI | SI | P6C4 | FASTA |
| 7G8 | ERS686280 | Brazil | SI | SI | P6C4 | FASTA |
| GB4 | ERS3566948 | Ghana | NIH | CSHL | P6C4 | FASTA |
| 803 | ERS3566949 | Cambodia | NIH | CSHL | P6C4 | FASTA |
| 36F11 | ERS3119776 | 803xGB4 progeny | NIH | CSHL | P6C4 | FASTA |

A machine-readable version of this manifest is available at `https://github.com/mcveanlab/CortexJDK/blob/master/manuscript/manifest.pacbio.txt`.

#### S3.2 Library preparation

To facilitate draft reference genome construction, we obtained high molecular weight genomic DNA (HMW gDNA) for seven *P. falciparum* parasites: all six parental clones spanning the four experimental crosses (3D7, HB3, DD2, 7G8, GB4, 803) and one progeny clone from the 803xGB4 cross (36F11). Parasite lines for 3D7, HB3, DD2, and 7G8 were produced by the Kwiatkowski lab at the Wellcome Trust Sanger Institute (SI). Lines for 803, GB4, and 36F11 were produced by the National Institute of Allergy and Infectious Diseases at the NIH (NIH). All cultures were maintained under standard conditions(11).

We obtained $6-20$ $\mu$g of HMW gDNA from each haploid parasite culture. QC was performed with NanoDrop spectrophotometers (Thermo Fisher Scientific) to verify gDNA purity. 20 kbp insert SMRTbell libraries were generated per sample with a Blue-Pippin (Sage Science) size selection range of $10-50$ kbp.

For additional assembly polishing of the HB3, DD2, and 7G8 genomes, we generated short fragment libraries using 0.5 $\mu g$ of DNA and a PCR-free library construction method(12). These libraries were sequenced using on MiSeq Illumina instruments, generating 250 bp paired-end reads and mean fragment length of 500 bp.

PacBio RSII sequencing of each library took place at two facilities. 3D7, GB4, 803, and 36F11 were sequenced at Cold Spring Harbor Laboratory's Next Generation Genomics Shared Resource (CSHL). HB3, DD2, and 7G8 were sequenced at the Sanger Institute.

#### S3.3 *De novo* assembly

3D7, HB3, DD2, and 7G8 assemblies on the libraries described above have been previously reported and publicly released(13). Summarizing, raw reads were assembled using the HGAP2 software using default parameters and a genome size of 23.5 Mbp. The HGAP assemblies were further improved by removing contaminating sequences, short contigs, and overlapping contigs; merging contigs with substantial overlap length; and polishing using the 250 bp Illumina reads aligned to the assembly.

Remaining samples GB4, 803, and 36F11 were assembled with the HGAP3 software.

##### S3.4 Contaminant removal

To remove possible contaminants from all assemblies, we ran contigs through BLAST(14), excluding any contig with a match to an organism other than *P. falciparum* in the `nt` database (updated Oct 2017).

##### S3.5 Pseudochromosome contiguation and gene annotation

To reorient contigs to match canonical reference orientation, build scaffolds representing whole chromosomes, and annotate genes, we ran all assemblies through the Companion webserver(15). We set the `Reference organism` setting as `Plasmodium falciparum 3D7`. We specified that both *ab initio* gene finding and existing gene model transfer be performed, specifying the `Strain` option for the RATT(16) gene model transfer tool. All other settings remained as software defaults. Note that Companion produced scaffolds representing all 14 autosomes in the *P. falciparum* genome, but does not automatically recognize and circularize the mitochondria and apicoplast genomes. These linear contigs, along with contigs that could not be placed in an autosome, are concatenated in a separate contig named with a "_00" suffix. Hence, scaffolding by pseudochromosome contiguation results in 15 total scaffolds per assembly.

##### S3.6 Annotation of repetitive regions and accessory compartments

We annotated repetitive sequences by applying the RepeatMasker(17) software to each genome using the maximum sensitivity `-s` option and the `-species 'plasmodium falciparum'` argument. We annotated core and accessory regions of each genome with the (18) software, processing each chromosome across all parental genomes simultaneously.

##### S3.7 Quality assessment

We first estimated the overall quality of our assemblies by performing a scaffold-to-chromosome alignment of our proof-of-principle 3D7 draft to the finished reference genome of the same parasite(19) using MUMmer(20). The alignments are visualized as a multi-dotplot in Figure S4, an extension of a dot plot that depicts alignments as two dimensional matrices with target and query sequences on the $x$ and $y$ axes respectively(21). Most chromosomes are assembled completely, and the overwhelming majority of the assembly appears on-diagonal (indicating successful one-to-one reconstruction). Elements appearing off-diagonal could represent misassembly. However, note that most of these off-diagonal elements occur towards the extremes of each chromosome. Given that the reference genome was constructed with Sanger reads an order of magnitude shorter than the PacBio reads, it is possible some repetitive regions have been collapsed or misplaced, contributing to this nominal error rate.

We called variants between the two assemblies to quantify errors using MUMmer, finding $3,357$ SNPs, $11,620$ insertions, and $4,603$ deletions. Overall, the SNP, insertion, and deletion rates are exceedingly low: amounting to $19,580$ events in a 23 Mbp genome (0.17%). The insertion rate is much higher than that of deletions and SNPs, perhaps due to the dominant insertion error mode of the PacBio sequencing instrument. All chromosomes appear reasonably similar in performance. Based on these measurements of the error rate, we estimate the quality of the PacBio assembly of the 3D7 isolate to be approximately Q31[1], or less than one error per thousand bases.

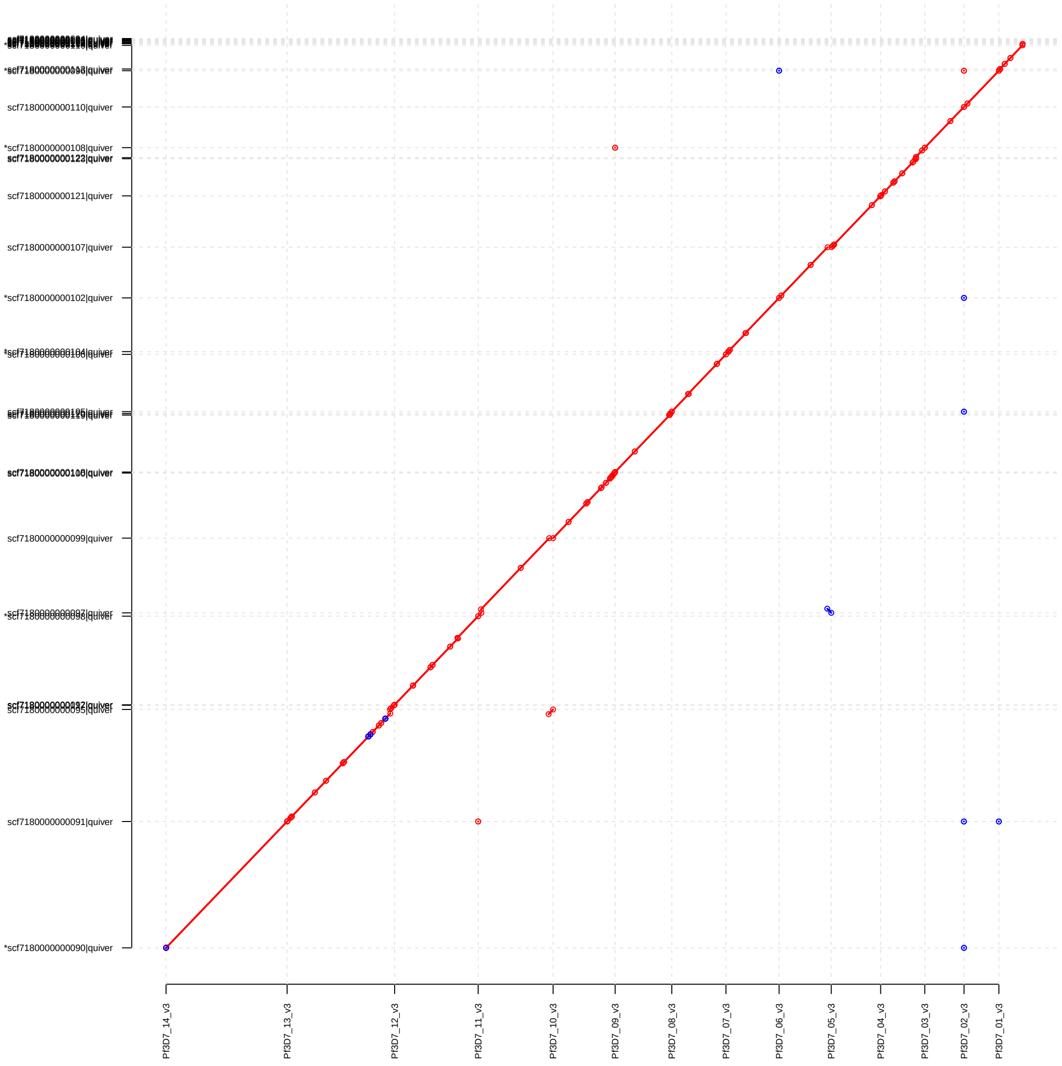

**Figure S4.** Alignment of contigs from the 3D7 draft to the finished reference assembly, with sequences from reference and draft assembly on the $x$ and $y$-axis, respectively. Each contig from the draft assembly is represented by a line segment terminated at either end with circles. Red denotes forward alignment of contigs to the reference, blue denotes reverse alignment.

As we did not have high-quality, finished genomes against which to compare the other assemblies, we devised a quality metric approximation based on $k$-mer counting. We used our graph processing software to find $k$-mers present in the PacBio assembly and absent in the Illumina

[1] $Q = -10\log_{10}(q) = -10\log_{10}((11,620 + 4,603 + 3,357)/23,332,831)$

**Table S3.** Metrics on PacBio sequencing and assemblies

| | 3D7 (ref) | 3D7 (draft) | HB3 | DD2 | 7G8 | GB4 | 803 | 36F11 |
|---|---|---|---|---|---|---|---|---|
| Assembler | – | HGAP2 | HGAP2 | HGAP2 | HGAP2 | HGAP3 | HGAP3 | HGAP3 |
| Contigs | – | 34 | 28 | 16 | 17 | 104 | 35 | 33 |
| Scaffolds[a] | 16 | 15 | 15 | 15 | 15 | 15 | 15 | 15 |
| Length (bp) | 23332839 | 23729641 | 22813863 | 22682439 | 22832395 | 27105702 | 23074973 | 23429077 |
| Length (autosomes) (bp) | 22671410 | 23194002 | 22671410 | 22641998 | 22775193 | 24613844 | 22772320 | 22989675 |
| Core genome (bp) | 20574502 | – | 21047328 | 21058329 | 21137184 | 21514228 | 21093156 | – |
| Accessory genome (bp) | 2096908 | – | 1624082 | 1583669 | 1638009 | 3099616 | 1679164 | – |
| Genes (autosomes) | 5561 | 5141 | 5130 | 5166 | 5219 | 5041 | 4501 | 4174 |
| Quality (MUMmer) | – | 31 | – | – | – | – | – | – |
| Quality (Graph) | – | 28 | 29 | 31 | 27 | 23 | 28 | 25 |

[a] In the draft reference sequences, the circular mitochondrial and apicoplast genomes are grouped together in a non-autosomal scaffold, resulting in one less scaffold than the canonical reference.

data for the same sample. We retained only one $k$-mer per variant event, discarding adjacent $k$-mers that tag the same event. We computed assembly quality using the number of retained $k$-mers as an estimate of the number of bases different between the PacBio and Illumina data.

We note that both measurements of assembly quality are pessimistic estimates. Our MUMmer-based estimates are predicated on the assumption that any differences between our 3D7 assembly and the canonical reference assembly indicate errors in our draft. In long, repetitive regions of the genome, this assumption may not be accurate as the Sanger reads used to assemble the reference were an order of magnitude shorter than the PacBio reads, potentially leading to collapsed repeats. Similarly for the $k$-mer counting estimate, while the PacBio RSII reads are known to have a much higher error rate than Illumina reads, it is not necessarily the case that all discrepancies between PacBio assemblies and Illumina reads should be adjudicated in favor of the latter.

Per-sample sequencing and final assembly details are presented in Table S3.

### S4 Data processing: short-read sequencing, alignment, and assembly

**Table** S4: Illumina metadata and download paths.

| cross | sample | source code | accession | fastq end1 ftp | fastq end2 ftp |
|---|---|---|---|---|---|
| 3D7xHB3 | PG0051-C | 3D7_Glasgow | ERR019061 | FASTQ (END1) | FASTQ (END2) |
| 3D7xHB3 | PG0052-C | HB3_Glasgow | ERR019054 | FASTQ (END1) | FASTQ (END2) |
| 3D7xHB3 | PG0053-C | XP3 | ERR019067 | FASTQ (END1) | FASTQ (END2) |
| 3D7xHB3 | PG0054-C | XP4 | ERR019062 | FASTQ (END1) | FASTQ (END2) |
| 3D7xHB3 | PG0055-C | XP5 | ERR019066 | FASTQ (END1) | FASTQ (END2) |
| 3D7xHB3 | PG0056-C | XP8 | ERR019068 | FASTQ (END1) | FASTQ (END2) |
| 3D7xHB3 | PG0057-C | XP9 | ERR019069 | FASTQ (END1) | FASTQ (END2) |
| 3D7xHB3 | PG0058-C | XP24 | ERR019063 | FASTQ (END1) | FASTQ (END2) |
| 3D7xHB3 | PG0060-C | XP52 | ERR019058 | FASTQ (END1) | FASTQ (END2) |
| 3D7xHB3 | PG0061-C | X2 | ERR019059 | FASTQ (END1) | FASTQ (END2) |
| 3D7xHB3 | PG0062-C | X4 | ERR019070 | FASTQ (END1) | FASTQ (END2) |
| 3D7xHB3 | PG0063-C | X5 | ERR019060 | FASTQ (END1) | FASTQ (END2) |
| 3D7xHB3 | PG0064-C | X6 | ERR019071 | FASTQ (END1) | FASTQ (END2) |
| 3D7xHB3 | PG0065-C | X10 | ERR019064 | FASTQ (END1) | FASTQ (END2) |
| 3D7xHB3 | PG0066-C | X11 | ERR019072 | FASTQ (END1) | FASTQ (END2) |
| 3D7xHB3 | PG0067-C | X12 | ERR019073 | FASTQ (END1) | FASTQ (END2) |
| 3D7xHB3 | PG0068-C | X30 | ERR019065 | FASTQ (END1) | FASTQ (END2) |
| 3D7xHB3 | PG0069-C | X33 | ERR019055 | FASTQ (END1) | FASTQ (END2) |
| 3D7xHB3 | PG0070-C | X35 | ERR019056 | FASTQ (END1) | FASTQ (END2) |
| 3D7xHB3 | PG0071-C | X39 | ERR019074 | FASTQ (END1) | FASTQ (END2) |
| HB3xDD2 | PG0004-CW | HB3_Ferdig | ERR012788 | FASTQ (END1) | FASTQ (END2) |
| HB3xDD2 | PG0008-CW | DD2_Ferdig | ERR012840 | FASTQ (END1) | FASTQ (END2) |
| HB3xDD2 | PG0015-C | B1SD | ERR019044 | FASTQ (END1) | FASTQ (END2) |
| HB3xDD2 | PG0016-C | QC13 | ERR012895 | FASTQ (END1) | FASTQ (END2) |
| HB3xDD2 | PG0017-C | QC01 | ERR019050 | FASTQ (END1) | FASTQ (END2) |
| HB3xDD2 | PG0018-C | B4R3 | ERR019042 | FASTQ (END1) | FASTQ (END2) |
| HB3xDD2 | PG0019-C | SC05 | ERR019051 | FASTQ (END1) | FASTQ (END2) |
| HB3xDD2 | PG0020-C | TC08 | ERR019052 | FASTQ (END1) | FASTQ (END2) |
| HB3xDD2 | PG0021-C | GC03 | ERR015447 | FASTQ (END1) | FASTQ (END2) |
| HB3xDD2 | PG0024-C | 3BD5 | ERR019053 | FASTQ (END1) | FASTQ (END2) |
| HB3xDD2 | PG0027-C | TC05 | ERR015450 | FASTQ (END1) | FASTQ (END2) |
| HB3xDD2 | PG0030-C | 7C188 | ERR019046 | FASTQ (END1) | FASTQ (END2) |
| HB3xDD2 | PG0031-C | 7C408 | ERR015458 | FASTQ (END1) | FASTQ (END2) |
| HB3xDD2 | PG0034-C | 7C3 | ERR019047 | FASTQ (END1) | FASTQ (END2) |
| HB3xDD2 | PG0037-C | 7C20 | ERR015451 | FASTQ (END1) | FASTQ (END2) |
| HB3xDD2 | PG0038-C | 7C111 | ERR015457 | FASTQ (END1) | FASTQ (END2) |
| HB3xDD2 | PG0039-C | 7C140 | ERR015454 | FASTQ (END1) | FASTQ (END2) |
| HB3xDD2 | PG0040-Cx | 7C159 | ERR107475 | FASTQ (END1) | FASTQ (END2) |
| HB3xDD2 | PG0041-C | 7C170 | ERR015446 | FASTQ (END1) | FASTQ (END2) |

| | | | | | |
|---|---|---|---|---|---|
| HB3xDD2 | PG0042-C | 7C183 | ERR015448 | FASTQ (END1) | FASTQ (END2) |
| HB3xDD2 | PG0043-C | 7C421 | ERR015459 | FASTQ (END1) | FASTQ (END2) |
| HB3xDD2 | PG0044-C | 7C424 | ERR019043 | FASTQ (END1) | FASTQ (END2) |
| HB3xDD2 | PG0045-C | QC23 | ERR012892 | FASTQ (END1) | FASTQ (END2) |
| HB3xDD2 | PG0046-Cx | 7C46 | ERR107476 | FASTQ (END1) | FASTQ (END2) |
| HB3xDD2 | PG0047-C | 7C126 | ERR015452 | FASTQ (END1) | FASTQ (END2) |
| HB3xDD2 | PG0048-C | 7C7 | ERR019049 | FASTQ (END1) | FASTQ (END2) |
| 7G8xGB4 | PG0077-C | JC3 | ERR027112 | FASTQ (END1) | FASTQ (END2) |
| 7G8xGB4 | PG0077-CW | JC3 | ERR045636 | FASTQ (END1) | FASTQ (END2) |
| 7G8xGB4 | PG0078-C | QF5 | ERR029092 | FASTQ (END1) | FASTQ (END2) |
| 7G8xGB4 | PG0078-CW | QF5 | ERR045638 | FASTQ (END1) | FASTQ (END2) |
| 7G8xGB4 | PG0079-C | JF6 | ERR027102 | FASTQ (END1) | FASTQ (END2) |
| 7G8xGB4 | PG0079-CW | JF6 | ERR045637 | FASTQ (END1) | FASTQ (END2) |
| 7G8xGB4 | PG0080-C | TF1 | ERR027103 | FASTQ (END1) | FASTQ (END2) |
| 7G8xGB4 | PG0081-CW | DEV_18_05_11 | ERR045633 | FASTQ (END1) | FASTQ (END2) |
| 7G8xGB4 | PG0082-C | WC4 | ERR029093 | FASTQ (END1) | FASTQ (END2) |
| 7G8xGB4 | PG0083-C | 7G8_NIH | ERR027099 | FASTQ (END1) | FASTQ (END2) |
| 7G8xGB4 | PG0084-C | GB4_NIH | ERR027100 | FASTQ (END1) | FASTQ (END2) |
| 7G8xGB4 | PG0087-C | JB8 | ERR029091 | FASTQ (END1) | FASTQ (END2) |
| 7G8xGB4 | PG0088-C | KH7 | ERR027111 | FASTQ (END1) | FASTQ (END2) |
| 7G8xGB4 | PG0090-C | KC2 | ERR027116 | FASTQ (END1) | FASTQ (END2) |
| 7G8xGB4 | PG0091-C | KA6 | ERR027117 | FASTQ (END1) | FASTQ (END2) |
| 7G8xGB4 | PG0093-C | XB3 | ERR029105 | FASTQ (END1) | FASTQ (END2) |
| 7G8xGB4 | PG0094-C | D2 | ERR027106 | FASTQ (END1) | FASTQ (END2) |
| 7G8xGB4 | PG0094-CW | D2_18_05_11 | ERR045632 | FASTQ (END1) | FASTQ (END2) |
| 7G8xGB4 | PG0095-C | NIC | ERR027107 | FASTQ (END1) | FASTQ (END2) |
| 7G8xGB4 | PG0096-C | NF10 | ERR027108 | FASTQ (END1) | FASTQ (END2) |
| 7G8xGB4 | PG0097-C | WF12 | ERR027109 | FASTQ (END1) | FASTQ (END2) |
| 7G8xGB4 | PG0098-C | DAN | ERR027110 | FASTQ (END1) | FASTQ (END2) |
| 7G8xGB4 | PG0099-C | JB12 | ERR029146 | FASTQ (END1) | FASTQ (END2) |
| 7G8xGB4 | PG0100-C | JE11 | ERR029404 | FASTQ (END1) | FASTQ (END2) |
| 7G8xGB4 | PG0101-C | KC5 | ERR029147 | FASTQ (END1) | FASTQ (END2) |
| 7G8xGB4 | PG0102-C | XF12 | ERR029143 | FASTQ (END1) | FASTQ (END2) |
| 7G8xGB4 | PG0102-CW | XF12_18_05_11 | ERR045635 | FASTQ (END1) | FASTQ (END2) |
| 7G8xGB4 | PG0104-C | KB8 | ERR029148 | FASTQ (END1) | FASTQ (END2) |
| 7G8xGB4 | PG0105-C | XD8 | ERR029144 | FASTQ (END1) | FASTQ (END2) |
| 7G8xGB4 | PG0105-CW | XD8_13_05_11 | ERR045628 | FASTQ (END1) | FASTQ (END2) |
| 7G8xGB4 | PG0107-C | JON | ERR029408 | FASTQ (END1) | FASTQ (END2) |
| 7G8xGB4 | PG0109-C | XG10 | ERR029405 | FASTQ (END1) | FASTQ (END2) |
| 7G8xGB4 | PG0110-C | LC12 | ERR171454 | FASTQ (END1) | FASTQ (END2) |
| 7G8xGB4 | PG0111-C | JC9 | ERR029409 | FASTQ (END1) | FASTQ (END2) |
| 7G8xGB4 | PG0111-CW | JC9_18_05_11 | ERR045634 | FASTQ (END1) | FASTQ (END2) |
| 7G8xGB4 | PG0112-C | AUD | ERR029406 | FASTQ (END1) | FASTQ (END2) |
| 7G8xGB4 | PG0113-CW | JH6_12_05_11 | ERR045626 | FASTQ (END1) | FASTQ (END2) |

| | | | |
|---|---|---|---|
| 803xGB4 | PG0443-C | 803 | ERR570006 FASTQ (END1) FASTQ (END2) |
| 803xGB4 | PG0050-CX2 | GB4 | ERR570014 FASTQ (END1) FASTQ (END2) |
| 803xGB4 | PG0445-C | 11H5 | ERR570030 FASTQ (END1) FASTQ (END2) |
| 803xGB4 | PG0446-C | 36F11 | ERR570038 FASTQ (END1) FASTQ (END2) |
| 803xGB4 | PG0447-C | 36H9 | ERR656280 FASTQ (END1) FASTQ (END2) |
| 803xGB4 | PG0448-C | 37D9 | ERR656283 FASTQ (END1) FASTQ (END2) |
| 803xGB4 | PG0450-C | 39A4 | ERR656289 FASTQ (END1) FASTQ (END2) |
| 803xGB4 | PG0451-C | 39C5 | ERR656292 FASTQ (END1) FASTQ (END2) |
| 803xGB4 | PG0453-C | 43H3 | ERR656298 FASTQ (END1) FASTQ (END2) |
| 803xGB4 | PG0455-C | 46G9 | ERR570015 FASTQ (END1) FASTQ (END2) |
| 803xGB4 | PG0458-C | 61A12 | ERR570039 FASTQ (END1) FASTQ (END2) |
| 803xGB4 | PG0459-C | 61D3 | ERR656281 FASTQ (END1) FASTQ (END2) |
| 803xGB4 | PG0460-C | 61E6 | ERR656284 FASTQ (END1) FASTQ (END2) |
| 803xGB4 | PG0461-C | 61E8 | ERR656287 FASTQ (END1) FASTQ (END2) |
| 803xGB4 | PG0462-C | 71D6 | ERR656290 FASTQ (END1) FASTQ (END2) |
| 803xGB4 | PG0463-C | 76H10 | ERR656293 FASTQ (END1) FASTQ (END2) |
| 803xGB4 | PG0464-C | 85D3 | ERR656296 FASTQ (END1) FASTQ (END2) |
| 803xGB4 | PG0465-C | 87A11 | ERR656299 FASTQ (END1) FASTQ (END2) |
| 803xGB4 | PG0466-C | 87E7 | ERR570008 FASTQ (END1) FASTQ (END2) |
| 803xGB4 | PG0467-C | 40G11 | ERR570016 FASTQ (END1) FASTQ (END2) |
| 803xGB4 | PG0469-C | 50C5 | ERR570032 FASTQ (END1) FASTQ (END2) |
| 803xGB4 | PG0470-C | 24G11 | ERR570040 FASTQ (END1) FASTQ (END2) |
| 803xGB4 | PG0471-C | 36D5 | ERR656282 FASTQ (END1) FASTQ (END2) |
| 803xGB4 | PG0472-C | 36E5 | ERR656285 FASTQ (END1) FASTQ (END2) |
| 803xGB4 | PG0473-C | 38G5 | ERR656288 FASTQ (END1) FASTQ (END2) |
| 803xGB4 | PG0474-C | 39C3 | ERR656291 FASTQ (END1) FASTQ (END2) |
| 803xGB4 | PG0475-C | 4E8 | ERR656294 FASTQ (END1) FASTQ (END2) |
| 803xGB4 | PG0476-C | 34F5 | ERR656297 FASTQ (END1) FASTQ (END2) |
| 803xGB4 | PG0492-C | 34B1 | ERR905451 FASTQ (END1) FASTQ (END2) |
| 803xGB4 | PG0494-C | 35C2 | ERR905453 FASTQ (END1) FASTQ (END2) |
| 803xGB4 | PG0495-C | 38A6 | ERR905454 FASTQ (END1) FASTQ (END2) |
| 803xGB4 | PG0496-C | 38E11 | ERR905455 FASTQ (END1) FASTQ (END2) |
| 803xGB4 | PG0498-C | 39G5 | ERR905457 FASTQ (END1) FASTQ (END2) |
| 803xGB4 | PG0499-C | 39H5 | ERR905458 FASTQ (END1) FASTQ (END2) |
| 803xGB4 | PG0500-C | 40A6 | ERR905459 FASTQ (END1) FASTQ (END2) |
| 803xGB4 | PG0501-C | 40B12 | ERR905460 FASTQ (END1) FASTQ (END2) |
| 803xGB4 | PG0502-C | 40F4 | ERR905461 FASTQ (END1) FASTQ (END2) |
| 803xGB4 | PG0503-C | 40G2 | ERR905462 FASTQ (END1) FASTQ (END2) |
| 803xGB4 | PG0504-C | 43E5 | ERR905463 FASTQ (END1) FASTQ (END2) |
| 803xGB4 | PG0505-C | 44D4 | ERR905464 FASTQ (END1) FASTQ (END2) |
| 803xGB4 | PG0508-C | 76H10-Tk13 | ERR905468 FASTQ (END1) FASTQ (END2) |
| 803xGB4 | PG0509-C | 85G7 | ERR905469 FASTQ (END1) FASTQ (END2) |
| 803xGB4 | PG0510-C | 88C9 | ERR905470 FASTQ (END1) FASTQ (END2) |
| 803xGB4 | PG0512-C | 37F12 | ERR905473 FASTQ (END1) FASTQ (END2) |

##### S4.1 Data availability

Raw reads for *P. falciparum* parents and progeny were provided to us by the MalariaGen project(22). ERA accession numbers and fastq download paths are provided in Table S4.

A machine-readable version of this manifest is available at https://github.com/mcveanlab/CortexJDK/blob/master/manuscript/manifest.illumina.txt.

##### S4.2 Library preparation

We obtained short-read, paired-end whole genome sequence data for parent and progeny clones of the 3D7xHB3(23), HB3xDD2(24), 7G8xGB4(25), and 803xGB4(26) crosses. All samples were sequenced on Illumina platforms between 2010 and 2014 using a PCR-free library preparation protocol intended to reduce coverage biases associated with AT-rich templates. Avoiding PCR during library construction also removes the issue of replication errors that occur in early cycles being propagated to all subsequent copies, thus masquerading as DNMs. Across all four crosses, and excluding parasite clones, we obtained 127 samples (8 parents, 119 progeny). The data is summarized in Table S5.

##### S4.3 Alignment

**Table S5.** Summary of sequencing data for four *P. falciparum* crosses

| | 3D7xHB3 | HB3xDD2 | 7G8xGB4 | 803xGB4 |
|---|---|---|---|---|
| Parents | 2 | 2 | 2 | 2 |
| Progeny | 18 | 24 | 35 | 42 |
| Read length (bp) | 76 | 76 | 76 | 100 |
| Insert size (bp) | $300 \pm 29$ | $253 \pm 48$ | $293 \pm 21$ | $222 \pm 10$ |
| Coverage | $99 \pm 38$ | $121 \pm 91$ | $110 \pm 40$ | $205 \pm 106$ |
| Platform | Illumina GAII | Illumina GAII | Illumina GAII | Illumina HiSeq 2000 |
| Sequencing date | 2010 | 2009-2012 | 2010-2011 | 2014 |

We obtained the *P. falciparum* reference sequence(19) obtained from PlasmoDB(27) (build 32.0, April 2017). For reference-based analyses, we aligned all reads with `bwa mem` to this reference using default settings. We marked duplicate reads using the `MarkDuplicates` tool in the Picard suite, and applied the GATK base quality score recalibrator (BQSR) to improve the accuracy of the base quality scores(28). We computed coverage metrics with the GATK's `DepthOfCoverage` tool, and insert size metrics with Picard's `CollectInsertSizeMetrics` tool[2]. A summary of the sample data is presented in Table S5.

##### S4.4 Reference-based variant calls

For later comparison to our de Bruijn graph-based DNM calls, we applied two software packages to the 20 samples in the 3D7xHB3 cross. For SNVs and small indels, we applied the GATK's

[2] http://broadinstitute.github.io/picard/

HaplotypeCaller(3), specifying a ploidy of 1 and leaving all other parameters as software defaults. For larger and more complex structural variants (insertions, deletions, tandem duplications, inversions, and translocations), we applied the Delly2 software(4). No additional filtering was applied to these callsets in order to preserve maximum sensitivity for later comparison to our *k*-mer-counting analyses.

##### S4.5 *De novo* assembly

We constructed multi-color linked de Bruijn graphs (LdBGs) for all 127 samples using McCortex(6) with a four-step pipeline. First, to build raw (un-error-cleaned) graphs, we applied the `build` command with a *k*-mer size parameter of 47 bp for all samples. Next, we ran the `clean` command with default settings to produce error-cleaned graphs. Substantial coverage variation in the *P. falciparum* samples caused the default error-cleaning process to erroneously remove true genomic *k*-mers. Unaddressed, this lead to gapped assemblies with unusually short contigs, as *k*-mers necessary for haplotype threading were not present in the graph. We implemented a repair module (`RecoverExcludedKmers`) in Corticall, recovering *k*-mers that were present in the raw graph, absent in the clean graph, and present in the canonical reference or either parental PacBio draft assembly for the cross. We ran the McCortex `inferedges` tool on the error-recovered graph, ensuring adjacencies between records in the graph were properly recorded. Finally, the parents, children, PacBio draft references, and reference genome were joined into a single multi-color graph to facilitate downstream processing using the Corticall `Join` command (a low-memory equivalent to McCortex's `join` command). All graphs were output in sorted order to facilitate random access over the dataset in Corticall.

We constructed three link annotation sets for each sample graph using the McCortex `thread` command: one for each parental PacBio draft and one for a sample's paired end reads. For PacBio assemblies and reads, we constructed links using the two-way gap-filling option. For paired-end reads, we further improved these links by re-threading links using `thread`'s paired-end mode and minimum (maximum) fragment size parameters 0 bp (1000 bp). We then filtered out low-coverage links likely arising from sequencing errors according to McCortex's documentation.

#### S5 Simulation of progeny genomes

We simulated progeny genomes from a synthetic crossing of HB3 and DD2 parasites with known allelic recombinations and DNMs using the Corticall module `SimulateHaploidChild`. Our simulations output a VCF describing recombination operations and other permutations to the appropriate reference sequences. The newly generated progeny reference genomes were then used to simulate reads for *de novo* assembly and DNM calling.

##### S5.1 Homologous recombination

We began with the pseudochromosome contiguated assemblies of HB3 and DD2 from section S3.5, for which homologous chromosomes suitable for recombination were easily paired. Crossover number $c$ was taken to be Poisson-distributed, parameterized with values as determined in Miles *et al.* 2016(29) and shown in Equation 1 with rate parameter $\mu = \lambda d$, average crossover rate $\lambda = 0.0135\ cM^{-1}$ and map length $d(\ell) = 0.74\ell - 0.11$, where $\ell$ is chromosome length. Map length and crossover probability distributions are shown in Figure S5.

$$P\{c|\ell\} = \frac{\mu^c}{c!}e^{-\mu} = \frac{(\lambda d(\ell))^c}{c!}e^{-\lambda d(\ell)} \tag{1}$$

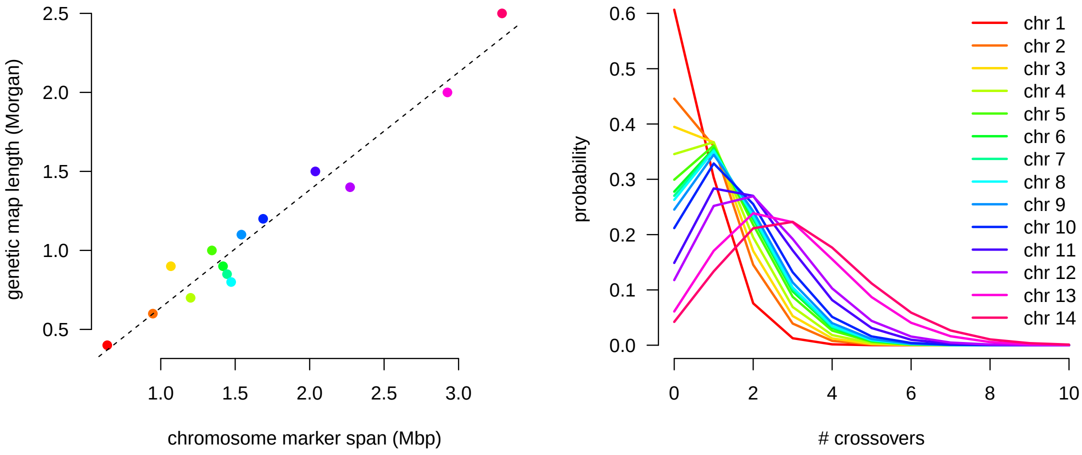

**Figure S5.** Simulated map lengths per chromosome (left) and crossover probability distribution per chromosome (right).

#### S5.2 Non-homologous allelic recombination

We simulated non-homologous allelic recombination (NAHR) events between members of the *var* gene family annotated in the HB3 and DD2 draft genome assemblies. While true NAHR events appear to be mediated by microhomology and possibly 5′ upstream promoter sequence, we deemed random pairings sufficient for our needs. For each simulated progeny, we chose at random two *var* genes from each genome, computed exact homology maps at $k = 21$ between the two sequences, and simulated a random number (between 1 and 6) of crossover events between the two sequences constrained to the homologous regions. We emitted two VCF entries for each NAHR event, the first replacing one *var* sequence with our new sequence, the other removing the second *var* gene from the progeny reference.

#### S5.3 Other variants

We simulated SNVs, short indels and inversions, multi-nucleotide polymorphisms. For the non-point mutations, we simulated a range of allele sizes up to $1,000$ bp. To evaluate our capabilities in expansions or contractions of repetitive sequence, we scanned the parental reference assemblies for series of repeating sequence units up to length 6 bp, adding or removing units up to the maximum number of units present in the repeat. All events were added to the VCF used for draft reference sequence permutation.

#### S5.4 Read simulation

Reads were simulated using 76 bp reads, insert size of 250 bp, a mean read depth of $120x$, and per-base error rate of 0.5%. All processing steps proceeded identically to the procedure specified in section S4.5.

#### S5.5 Evaluation

To evaluate performance of our mutation caller, we developed the `EvaluateCalls` module in Corticall. To overcome issues regarding different (but effectively equivalent) descriptions of alleles between the simulation VCF and the Corticall output VCF, we reused the reference permutation concept from S1.2. We permuted the indicated draft reference genomes with the simulated and called VCFs and evaluated local sequence equivalence between the two callsets. We emitted precision and recall metrics for each variant, and additional information regarding variants that had been incompletely assembled.

[2] Heng Li. Aligning sequence reads, clone sequences and assembly contigs with BWA-MEM. *arXiv.org*, page arXiv:1303.3997, March 2013.

[3] Ryan Poplin, Valentin Ruano Rubio, Mark A DePristo, Tim J Fennell, Mauricio O Carneiro, Geraldine A Van der Auwera, David E Kling, Laura D Gauthier, Ami Levy-Moonshine, David Roazen, Khalid Shakir, Joel Thibault, Sheila Chandran, Chris Whelan, Monkol Lek, Stacey Gabriel, Mark J Daly, Benjamin Neale, Daniel G MacArthur, and Eric Banks. Scaling accurate genetic variant discovery to tens of thousands of samples. *bioRxiv*, page 201178, November 2017.

[4] Tobias Rausch, Thomas Zichner, Andreas Schlattl, Adrian M Stütz, Vladimir Benes, and Jan O Korbel. DELLY: structural variant discovery by integrated paired-end and split-read analysis. 28(18):i333–i339, September 2012.

[5] Zamin Iqbal, Mario Caccamo, Isaac Turner, Paul Flicek, and Gil McVean. De novo assembly and genotyping of variants using colored de Bruijn graphs. *Nature genetics*, 44(2):226–232, February 2012.

[6] Isaac Turner, Kiran V Garimella, Zamin Iqbal, and Gil McVean. Integrating long-range connectivity information into de Bruijn graphs. *bioRxiv*, page 147777, June 2017.

[7] S F Altschul, W Gish, W Miller, E W Myers, and D J Lipman. Basic local alignment search tool. *Journal of molecular biology*, 215(3):403–410, October 1990.

[8] Ella K Chase Donald S Chen Philip Awadalla Karen P Day Gil McVean Martine M Zilversmit. Hypervariable antigen genes in malaria have ancient roots. *BMC Evolutionary Biology*, 13:110, 2013.

[9] Richard Durbin. *Biological Sequence Analysis*. Probabilistic Models of Proteins and Nucleic Acids. Cambridge University Press, April 1998.

[15] Sascha Steinbiss, Fatima Silva-Franco, Brian Brunk, Bernardo Foth, Christiane Hertz-Fowler, Matthew Berriman, and Thomas D Otto. Companion: a web server for annotation and analysis of parasite genomes. *Nucleic acids research*, 44(W1):W29–W34, July 2016.

[16] Thomas D Otto, Gary P Dillon, Wim S Degrave, and Matthew Berriman. RATT: Rapid Annotation Transfer Tool. *Nucleic acids research*, 39(9):e57–e57, May 2011.

[17] AFA Smit, R Hubley, and P Green. *RepeatMasker Open-4.0*, 2013.

[18] Egon A Ozer, Jonathan P Allen, and Alan R Hauser. Characterization of the core and accessory genomes of Pseudomonas aeruginosa using bioinformatic tools Spine and AGEnt. *BMC genomics*, 15(1):737, December 2014.

[19] Malcolm J Gardner, Neil Hall, Eula Fung, Owen White, Matthew Berriman, Richard W Hyman, Jane M Carlton, Arnab Pain, Karen E Nelson, Sharen Bowman, Ian T Paulsen, Keith James, Jonathan A Eisen, Kim Rutherford, Steven L Salzberg, Alister Craig, Sue Kyes, Man-Suen Chan, Vishvanath Nene, Shamira J Shallom, Bernard Suh, Jeremy Peterson, Sam Angiuoli, Mihaela Pertea, Jonathan Allen, Jeremy Selengut, Daniel Haft, Michael W Mather, Akhil B Vaidya, David M A Martin, Alan H Fairlamb, Martin J Fraunholz, David S Roos, Stuart A Ralph, Geoffrey I McFadden, Leda M Cummings, G Mani Subramanian, Chris Mungall, J Craig Venter, Daniel J Carucci, Stephen L Hoffman, Chris Newbold, Ronald W Davis, Claire M Fraser, and Bart Barrell. Genome sequence of the human malaria parasite Plasmodium falciparum. *Nature*, 419(6906):498–511, October 2002.

[20] Stefan Kurtz, Adam Phillippy, Arthur L Delcher, Michael Smoot, Martin Shumway, Corina Antonescu, and Steven L Salzberg. Versatile and open software for comparing large genomes. *Genome Biology*, 5(2):R12, January 2004.

[21] Adrian J Gibbs and George A Mcintyre. The Diagram, a Method for Comparing Sequences. *European Journal of Biochemistry*, 16(1):1–11, September 1970.

[22] Alistair Miles, Zamin Iqbal, Paul Vauterin, Richard Pearson, Susana Campino, Michel Theron, Kelda Gould, Daniel Mead, Eleanor Drury, John O'Brien, Valentin Ruano Rubio, Bronwyn MacInnis, Jonathan Mwangi, Upeka Samarakoon, Lisa Ranford-Cartwright, Michael Ferdig, Karen Hayton, Xinzhuan Su, Thomas Wellems, Julian Rayner, Gil McVean, and Dominic Kwiatkowski. Genome variation and meiotic recombination in Plasmodium falciparum: insights from deep sequencing of genetic crosses. *bioRxiv*, page 024182, August 2015.

[23] D Walliker, I Quakyi, T Wellems, T McCutchan, A Szarfman, W London, L Corcoran, T Burkot, and R Carter. Genetic analysis of the human malaria parasite Plasmodium falciparum. 236(4809):1661–1666, June 1987.

[24] T E Wellems, L J Panton, I Y Gluzman, V E do Rosario, R W Gwadz, A Walker-Jonah, and D J Krogstad. Chloroquine resistance not linked to mdr-like genes in a Plasmodium falciparum cross. *Nature*, 345(6272):253–255, May 1990.

[25] Karen Hayton, Deepak Gaur, Anna Liu, Jonathan Takahashi, Bruce Henschen, Subhash Singh, Lynn Lambert, Tetsuya Furuya, Rachel Bouttenot, Michelle Doll, Fatima Nawaz, Jianbing Mu, Lubin Jiang, Louis H Miller, and Thomas E Wellems. Erythrocyte Binding Protein PfRH5 Polymorphisms Determine Species-Specific Pathways of Plasmodium falciparum Invasion. *Cell Host & Microbe*, 4(1):40–51, January 2008.

[26] Juliana M Sá, Sarah R Kaslow, Michael A Krause, Viviana A Melendez-Muniz, Rebecca E Salzman, Whitney A Kite, Min Zhang, Roberto R Moraes Barros, Jianbing Mu, Paul K Han, J Patrick Mershon, Christine E Figan, Ramoncito L Caleon, Rifat S Rahman, Tyler J Gibson, Chanaki Amaratunga, Erika P Nishiguchi, Kimberly F Breglio, Theresa M Engels, Soundarapandian Velmurugan, Stacy Ricklefs, Judith Straimer, Nina F Gnadig, Bingbing Deng, Anna Liu, Ababacar Diouf, Kazutoyo Miura, Gregory S Tullo, Richard T Eastman, Sumana Chakravarty, Eric R James, Kenneth Udenze, Suzanne Li, Daniel E Sturdevant, Robert W Gwadz, Stephen F Porcella, Carole A Long, David A Fidock, Marvin L Thomas, Michael P Fay, B Kim Lee Sim, Stephen L Hoffman, John H Adams, Rick M Fairhurst, Xin-zhuan Su, and Thomas E Wellems. Artemisinin resistance phenotypes and K13 inheritance in a Plasmodium falciparum cross and Aotus model. *Proceedings of the National Academy of Sciences*, pages 1–6, November 2018.

[27] C Aurrecoechea, J Brestelli, B P Brunk, J Dommer, S Fischer, B Gajria, X Gao, A Gingle, G Grant, O S Harb, M Heiges, F Innamorato, J Iodice, J C Kissinger, E Kraemer, W Li, J A Miller, V Nayak, C Pennington, D F Pinney, D S Roos, C Ross, C J Stoeckert, C Treatman, and H Wang. PlasmoDB: a functional genomic database for malaria parasites. *Nucleic acids research*, 37(Database):D539–D543, January 2009.

[28] Mark A DePristo, Eric Banks, Ryan Poplin, Kiran V Garimella, Jared R Maguire, Christopher Hartl, Anthony A Philippakis, Guillermo del Angel, Manuel A Rivas, Matt Hanna, Aaron McKenna, Tim J Fennell, Andrew M Kernytsky, Andrey Y Sivachenko, Kristian Cibulskis, Stacey B Gabriel, David Altshuler, and Mark J Daly. A framework for variation discovery and genotyping using next-generation DNA sequencing data. *Nature genetics*, 43(5):491–498, May 2011.

[29] Alistair Miles, Zamin Iqbal, Paul Vauterin, Richard Pearson, Susana Campino, Michel Theron, Kelda Gould, Daniel Mead, Eleanor Drury, John O'Brien, Valentin Ruano Rubio, Bronwyn MacInnis, Jonathan Mwangi, Upeka Samarakoon, Lisa Ranford-Cartwright, Michael Ferdig, Karen Hayton, Xin-Zhuan Su, Thomas Wellems, Julian Rayner, Gil McVean, and Dominic Kwiatkowski. Indels, structural variation, and recombination drive genomic diversity in Plasmodium falciparum. *Genome Research*, 26(9):1288–1299, September 2016.
